# Prime editing of the β_1_ adrenoceptor in the brain reprograms mouse behavior

**DOI:** 10.1101/2023.05.19.541410

**Authors:** Desirée Böck, Lisa Tidecks, Maria Wilhelm, Yanik Weber, Eleonora Ioannidi, Jonas Mumenthaler, Tanja Rothgangl, Lukas Schmidheini, Sharan Janjuha, Tommaso Patriarchi, Gerald Schwank

## Abstract

Prime editing is a highly versatile genome editing technology that holds great potential for treating genetic diseases^1, 2^. While *in vivo* prime editing has recently been conducted in the brain, liver, heart, and retina^3–6^, application of this technology to modulate neural circuits in the brain has not been reported yet. Here, we employ adeno-associated viral vectors to deliver optimized intein-split prime editors into the brain of mice. Delivery into newborn pups via intracerebroventricular injection resulted in up to 44.0% editing at the *Dnmt1* locus in the cortex (on average 34.8±9.8% after 6 months). In addition, we obtained up to 28.1% editing at the *Adrb1* locus in the cortex (on average 14.7±11.6% after 6 months). The introduced *Adrb1*^A187V^ mutation is a naturally occurring variant of the β1-adrenergic receptor, which has previously been linked to increased activity and natural short sleep^7^. Similarly, we observed an increase in the activity and exploratory behavior of treated animals. This study demonstrates the potential of prime editing for treating genetic diseases in the central nervous system and for reprogramming molecular pathways that modulate animal behavior.

## Introduction

Prime editing is a versatile genome editing technology that enables the installation of all small-sized genetic changes, including transversion/transition mutations, insertions, and deletions^1^. Prime editors (PEs) consist of an *Sp*Cas9 nickase (H840A) fused to an engineered reverse transcriptase (RT) derived from the Moloney murine leukemia virus (M-MLV; hereafter referred to as PE2)^1^. The *Sp*Cas9-RT fusion protein further complexes with a prime editing guide RNA (pegRNA), which consists of a primer binding site (PBS) and an edit-containing programmable RT template (RTT) fused to the 3’ end of the guide RNA. In contrast to classical Cas9 nucleases, prime editing does not require the induction of error-prone DNA double-strand breaks and does not rely on homology-directed repair (HDR) to achieve accurate editing. Therefore, it also facilitates precise correction of mutations in non-dividing cells such as hepatocytes and retinal cells^3–5^.

Despite its potential for therapy, the efficiency of prime editing is still limited compared to Cas9 nucleases and base editors^1, 4, 8–10^. Several recent studies have attempted to improve the performance of prime editing by i) co-delivery of nicking sgRNAs (ngRNAs) that cut the non-edited strand simultaneously to the PE (PE3 ngRNAs) or after resolution of the edited strand (PE3b ngRNAs)^1, 2, 4, 11^, ii) adding RNA-stabilizing pseudoknot structures to the 3’ end of the pegRNA (hereafter referred to as epegRNAs)^11^, iii) using a codon-optimized PE variant that harbors additional mutations in the Cas9 nickase domain (R221K, N394K) and an adjusted linker/NLS design (PEmax)^2^, and iv) inhibiting DNA-mismatch repair (MMR) via co-expression of a dominant-negative MutL homolog 1 (dnMLH1)^2^. While these methods have been demonstrated to improve prime editing *in vitro*, their effect on *in vivo* prime editing efficiency is not well explored.

In this study, we developed a prime editing approach in the mouse brain using optimized pegRNA designs and expression vectors for adeno-associated virus (AAV) delivery. In addition to targeting *Dnmt1*, we introduced a behavior-affecting mutation in *Adrb1*, which encodes for the β1-adrenoceptor (β1-AR). β-ARs are activated by the neurotransmitter norepinephrine (NE; Extended data fig. 1a)^12^, and involved in a wide range of brain functions, including sleep/wake regulation^13^, memory^14^, and regulation of stress-and anxiety-induced responses^15, 16^. β1-ARs are particularly important for physiological functions of the sympathetic nervous system^17, 18^, with the naturally occurring variant *Adrb1*^A187V^ being linked to increased activity and short sleep in humans and mice^7^. Moreover, *Adrb1*^A187V^ has recently been shown to improve sleep quality and to ameliorate tau accumulation in a mouse model of tauopathy^19^, showing that effects mediated by β1-ARs can have various implications on health and disease. Using our optimized prime editing approach, we were able to introduce this mutation in various brain areas, leading to a change in animal behavior towards a heightened activity state.

**Figure 1:**
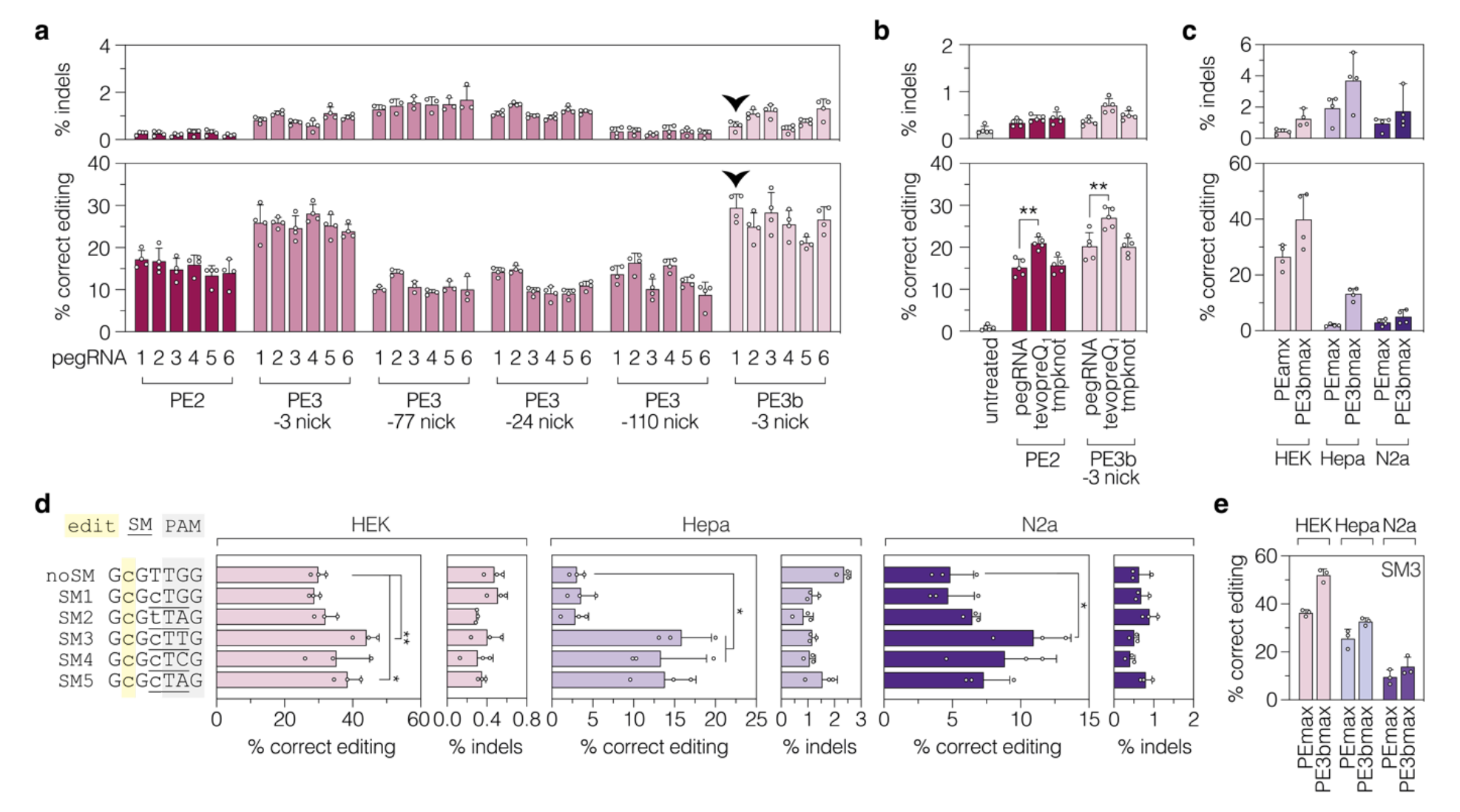
Optimization of pegRNAs and ngRNAs at the *Adbr1* locus. (**a**) Editing and indel rates of six pegRNAs alone (PE2) or combined with different PE3 ngRNAs or a PE3b ngRNA in HEK Adrb1 reporter cells. The location of the second nick relative to the installed edit is indicated. The black arrowhead labels pegRNA1, which was used for subsequent experiments. (**b**) Editing and indel rates of tevopreQ_1_- and tmpknot-epegRNA1 in HEK Adrb1 reporter cells. Data are displayed as means±s.d. of at least three independent experiments. nt, nucleotides. (**c**) Editing and indel ratets of epegRNA1 in combination with a PE3b ngRNA in HEK, Hepa, N2a Adrb1 reporter cells. (**d**) Editing and indel rates of epegRNAs encoding for SMs in HEK, Hepa, N2a Adrb1 reporter cells. The position of the edit (yellow), PAM sequence (gray), and SMs (underlined) are indicated. (**e**) Editing and indel rates of epegRNA-SM^CTT^ without (PEmax) and with (PE3bmax) a ngRNA. Data are displayed as means±s.d. of at least three independent experiments and were analyzed using a two-tailed Student’s t-test with Welch’s correction (*\*P<0.05*; *\*\*P<0.005*). If not indicated, differences were not statistically significant (*P>0.05*).

## Results

### Installation of the *Adrb1*^A187V^ mutation via prime editing in cell lines

To develop a prime editing approach for efficient installation of the A187V mutation in the *Adrb1* gene, we first tested different pegRNA designs at the endogenous locus in murine cell lines. We designed six pegRNAs encoding for the respective C-to-T edit with varying lengths of the RTT (11, 13, or 15 nucleotides [nt]) and PBS (10 or 13nt; Extended data fig. 1b). Vectors expressing the pegRNAs were first co-delivered with a PE2-expressing plasmid into murine Hepa1-6 and Neuro2a cell lines (hereafter referred to as Hepa and N2a). However, since deep sequencing of the target locus revealed low editing rates with all tested pegRNAs (< 2%; Extended data fig. 1c), potentially because the *Adrb1* is inaccessible in these cell lines (Extended data fig. 1d,e), we next generated HEK cells where the targeted *Adrb1* region is integrated into the genome using the PiggyBac transposon system^20^ (Extended data fig. 1f). Transfection of PE2 together with the different pegRNAs, either alone or in combination with different ngRNAs, identified pegRNA1 with a 10-nt PBS and 11-nt RTT as the most efficient pegRNA in installing the *Adrb1*^A187V^ mutation without inducing high rates of indels (Fig. 1a; extended data fig. 1b).

Structural 3’ modifications that protect pegRNAs from exonucleases have previously been reported to enhance editing efficiencies^11^. Therefore, we next tested whether fusing the stabilizing structural motifs tevopreQ_1_ and tmpknot to the 3’ end further enhances activity of pegRNA1. While adding the tmpknot motif did not have an impact on editing rates, adding the tevopreQ_1_ motif resulted in a 1.4-fold increase in pegRNA1 activity (Fig. 1b). Notably, we also tested if adding stabilizing motifs to the 3’ end of ngRNAs increased editing efficiency, but did not observe significant changes at various loci (Extended data fig. 2). Finally, we assessed whether exchanging the scaffold of the pegRNA with an optimized sgRNA scaffold harboring an extended duplex length and a base substitution in the poly-thymidine stretch further increases editing^21^. Although statistically not significant, the optimized scaffold led to higher activity of pegRNA1, with-and without the tevopreQ_1_ motif (Extended data fig. 3). Based on these results, we decided to select the tevopreQ_1_-modified pegRNA1 with the optimized scaffold (epegRNA1) for subsequent experiments.

**Figure 2:**
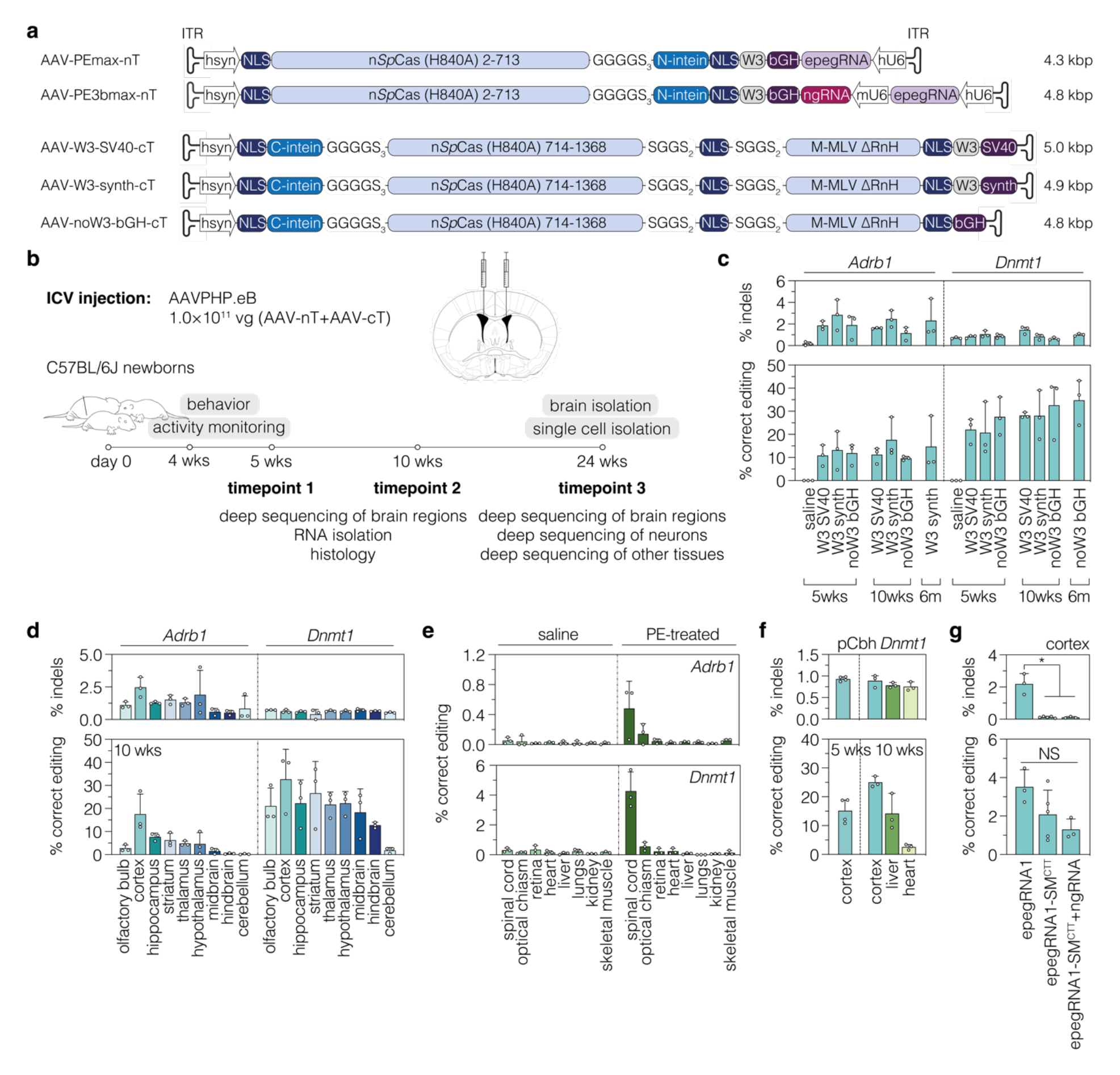
*In vivo* prime editing at the *Dnmt1* and *Adrb1* locus in the brain. (**a**) Schematic representation of AAV designs used *in vivo* and their corresponding lengths in kilobase pairs (kbp, including ITRs) for neuron-specific expression of PEmax or PE3bmax. Constructs are not depicted to scale. (**b**) Schematic representation of the experimental setup and timeline. (**c**) *In vivo* prime editing and indel rates of different AAV vector designs in mouse cortices at 5 weeks, 10 weeks and 6 months post-injection. (**d**) Editing and indel rates at the *Adrb1* (AAV-PE3bmax-nT and W3-synth-cT) and *Dnmt1* (AAV-PEmax-nT and noW3-bGH-cT) locus in different brain regions at 10 weeks post-injection. (**e**) Frequency of *Adrb1* and *Dnmt1* edits in various other tissues in saline-and phsyn-PE-treated animals at 6 months post injection. Animals were treated with the same AAV preparations as in (d). Skeletal muscle tissue was isolated from the quadriceps femoris. (**f**) Editing rates at the *Dnmt1* locus in mouse cortices (5 and 10 weeks post-injection), liver, and heart (10 weeks post-injection) after ICV injection. Animals were treated with PEmax-noW3-synth-nT and noW3-synth-cT under the control of the ubiquitous Cbh promoter^38^. (**g**) Comparison of editing and indel rates at the *Adrb1* locus for PEmax complexed with epegRNA1 or epegRNA1-SM^CTT^ (with and without a PE3b-ngRNA) at 5 weeks. Animals were treated with AAV-PEmax-nT and AAV-W3-synth-cT. Data are displayed as means±s.d. of at least three animals. Each data point represents one animal. ITR, inverted terminal repeat; nT/cT, N-/C-terminal PEmax AAV vector; phsyn, human synapsin promoter; NLS, nuclear localization signal; n*Sp*Cas9, *Sp*Cas9 nickase; M-MLV, Moloney Murine Leukemia virus; W3, woodchuck hepatitis virus post-transcriptional regulatory element; hU6/mU6, human/mouse U6 promoter; SV40, Simian virus 40; pA, polyA signal; synth, synthetic polyA signal; bGH, bovine growth hormone polyA signal; kbp, kilobasepairs; vg, vector genomes; wks, weeks; m, months; SM, silent mutation; pCbh, truncated chimeric CMV/chicken-β-actin hybrid promoter.

**Figure 3:**
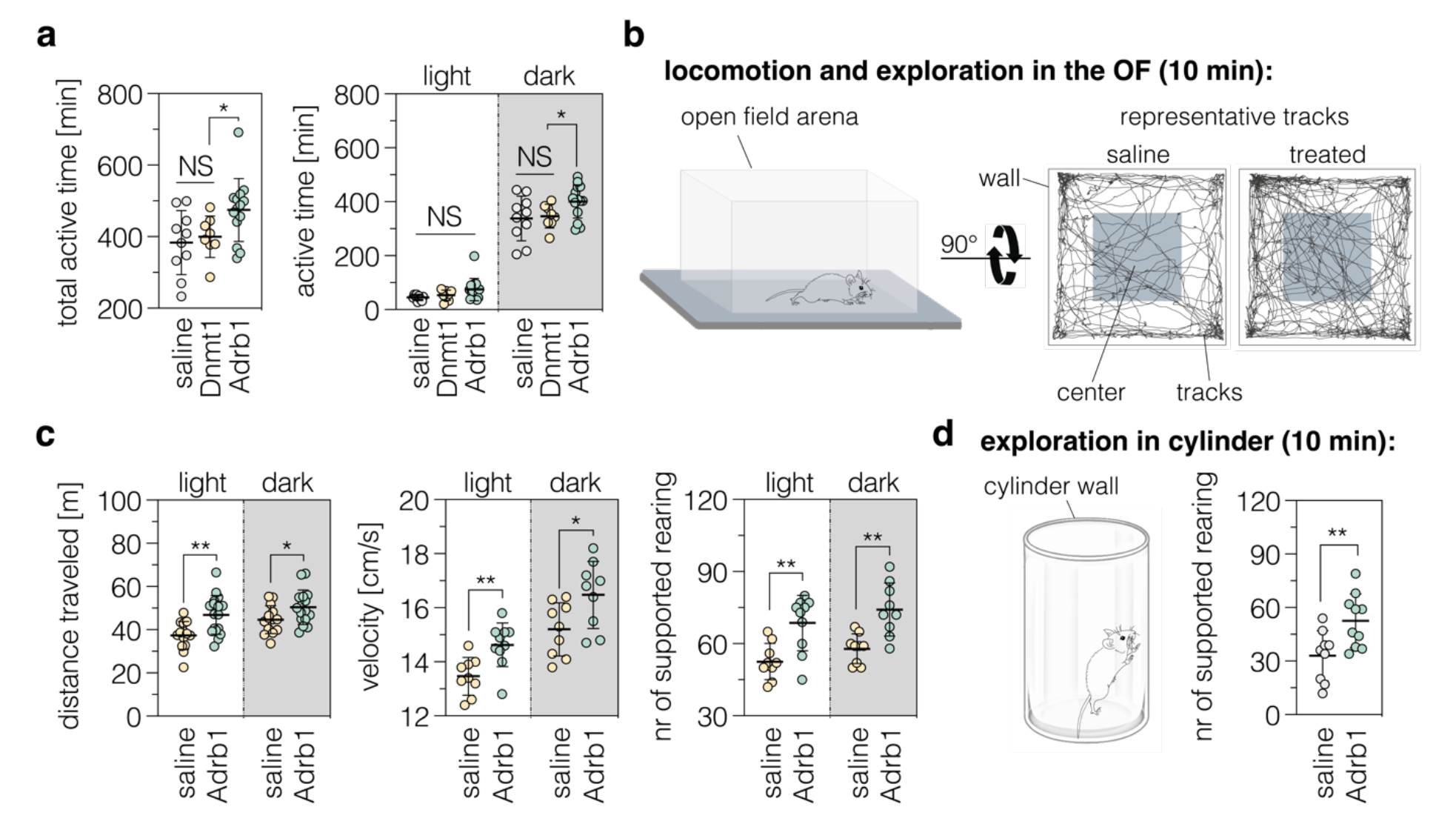
The *Adrb1*^A187V^ mutation increases locomotor activity and exploratory behavior in newborn mice. (**a**) General activity (expressed as total active time) in the home cage during the light and dark cycle (n=9-15 mice per group). (**b**) Schematic representation of the OF test and representative tracks of control and treated mice. (**c**) Locomotor activity, velocity, and supported rearing in the OF (n=9-15 mice per group). (**d**) Schematic representation and quantification of supported rearing in a cylinder (n=9-10 mice per group). The duration of each test is indicated in brackets (b,d). PE-treated and saline-injected mice were kept in a 12:12 light/dark cycle; areas highlighted in gray indicate the dark cycle. Data are displayed as means±s.d. and were analyzed using a two-tailed Student’s *t*-test with Welch’s correction (*\*P<0.05*; *\*\*P<0.005*). Each data point represents one animal. NS, not significant.

### Optimizing epegRNA1 for MMR proficient cells

DNA mismatch repair (MMR) has been shown to impede prime editing and promote undesired indel byproducts^2, 22^. This could be a limitation for *in vivo* prime editing in the brain, which contains MMR proficient cells^23^. Therefore, we next integrated the *Adrb1* reporter into MMR proficient Hepa-and N2a cells and assessed editing rates of epegRNA1 with PE2 or the optimized PE variant PEmax. In line with the hypothesis that MMR negatively affects prime editing, editing rates were significantly lower than in HEK cells (1.6±1.5% for PE2 and 2.0±0.4% for PEmax in Hepa cells; 0.4±0.1% for PE2 and 3.0±1.3% for PEmax in N2a cells; extended data fig. 4d), and inhibiting MMR via shRNA-mediated downregulation of *Mlh1* or *Msh2* or co-expression of dominant negative MLH1^2^ led to an increase in editing rates (up to 3.2-fold in Hepa-and 3.4-fold in N2a cells; extended data fig. 4a-d). Since MMR defects are a major driver for different cancers^24^, *in vivo* inhibition of MMR is likely detrimental, prompting us to assess if the inhibitory effect of MMR on prime editing could be circumvented by i) using the PE3b approach, or ii) modifying epegRNA1 to additionally introduce silent mutations (SMs) in the protospacer and PAM^2, 25^. Co-transfection of a PE3b ngRNA indeed led to a pronounced increase of editing rates in Hepa (6.8-fold) and N2a cells (1.7-fold; fig.1c). Likewise, three of the five epegRNA1 variants encoding for additional SMs led to substantially higher editing in Hepa (up to 5.2-fold) and N2a cells (up to 2.3-fold) in an MMR independent manner (Fig. 1d; extended data fig. 4e), with epegRNA1-SM^CTT^ showing highest editing efficiencies (HEK: 44.0±3.7%; Hepa: 15.9±3.6%; N2a: 11.0±2.7%). When combined with a PE3b ngRNA, editing rates of epegRNA1-SM^CTT^ could be even further enhanced (Fig. 1e). Notably, epegRNAs encoding for SMs that edit the PAM, and hence abolish re-targeting of the site after installation of the edit, also resulted in a substantial reduction of indel rates (Fig. 1c). Based on these results, we decided to use PEmax together with epegRNA1 and epegRNA1- SM^CTT^ in the PEmax or PE3b approach for subsequent *in vivo* experiments.

**Figure 4:**
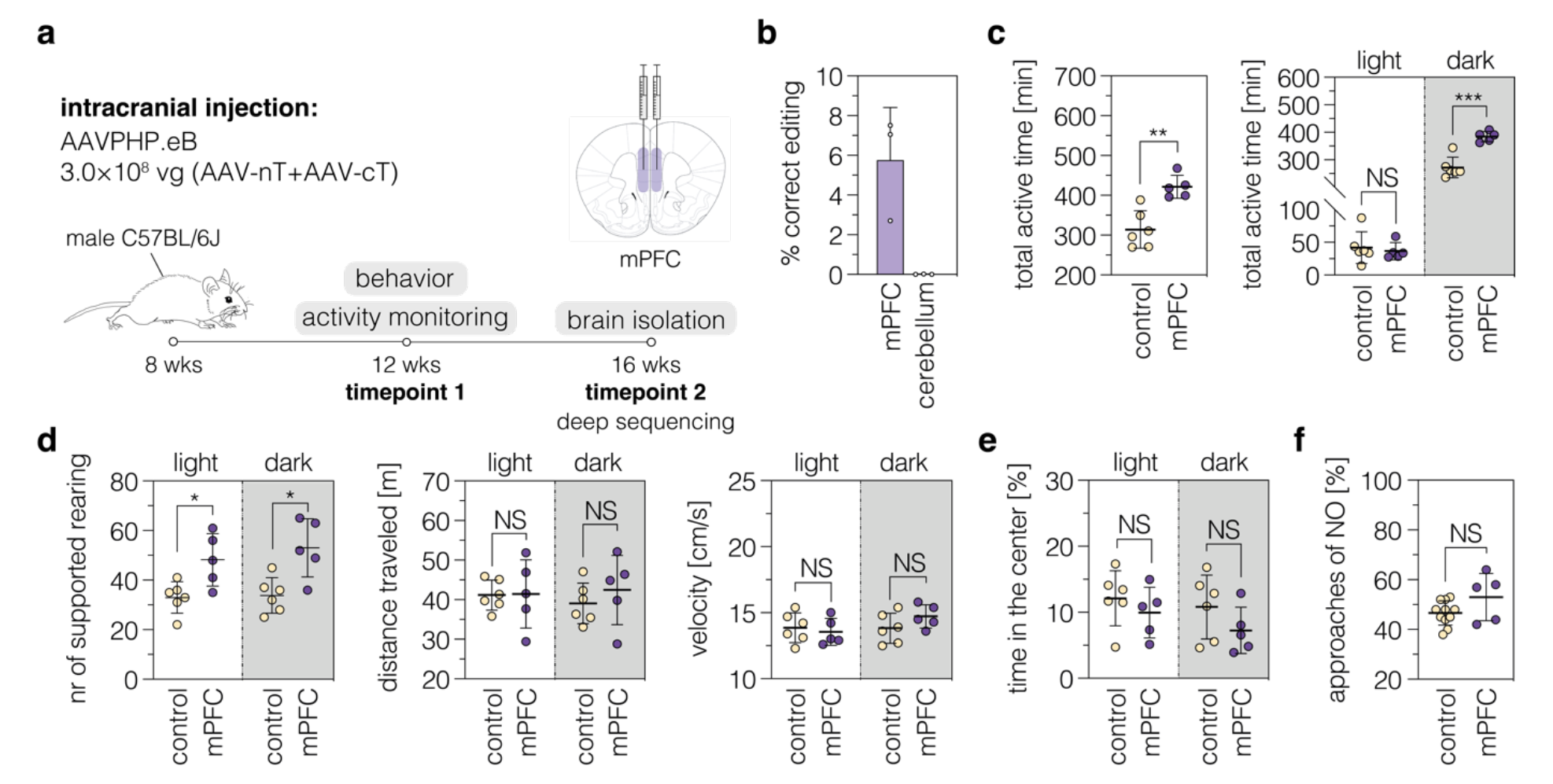
*Adrb1*^A187V^ in the mPFC increases general activity and exploration in adult mice. (**a**) Schematic depiction of the experimental setup used to introduce the *Adrb1*^A187V^ mutation into the mPFC of adult male mice. (**b**) Editing and indel rates in the mPFC and surrounding areas of adult mice after intracranial delivery of prime editing components. The cerebellum was used as a negative control. (**c**) Comparison of general activity in the home cage of controls and animals treated in the mPFC. (**d**) Supported rearings, locomotor activity, and velocity in the OF for treated (n=5 mice) and control mice (n=6 mice) in the light and dark cycle. Data are displayed as means±s.d. of at least 5 animals per group and were analyzed using a two-tailed Student’s t-test with Welch’s correction (*\*P<0.05*; *\*\*P<0.005; ***P<0.0005*). Each data point represents one animal.

### Generation of optimized intein-split PE variants for AAV-mediated delivery

Due to low immunogenicity, rare genomic integration, and the ability to efficiently infect neurons and astrocytes, AAVs are promising vectors for *in vivo* delivery of genome editors into the central nervous system (CNS)^26^. As PEs exceed the packaging limit of AAVs (∼5 kb including ITRs)^27^, we have employed the *Npu* intein-mediated protein trans-splicing system to split the PE into two parts for expression from two separate AAVs^5, 28–32^.

To further optimize intein-split PE vectors towards accommodating a variety of promoters and terminators for wide-range usage, we first generated nine PE variants split at different surface-exposed sites and assessed their activity on a GFP reporter and the *Adrb1* locus. Of the tested variants PE-p.713/714 was the most efficient, maintaining over 90% of the activity of the unsplit PE (Extended data fig. 5a-b). Next, we generated AAV expression cassettes where the N-and C-terminal fragments of PE-p.713/714 were cloned downstream of the neuron-specific human synapsin (hsyn) promoter^33^. While the vector encoding for the N-terminal PE fragment was small enough to additionally harbor expression cassettes for the epegRNA and ngRNA (Fig. 2a), the C-terminal fragment would have exceeded the AAV packaging limit (Extended data fig. 5d). Therefore, we generated C-terminal constructs where we removed the dispensable RnaseH domain from the RT^5, 34^ (Fig. 2a; extended data fig. 5c), and exchanged the commonly used W3-bGH terminator^35, 36^ with smaller terminators. These either lacked the W3 post-transcriptional regulatory element, or used shorter polyadenylation signals (SV40 or synthetic polyA instead of bGH polyA) (Fig. 2a; Extended data fig. 5d). *In vitro* comparison of the shorter terminators to the W3-bGH terminator revealed higher editing rates with the bGH terminator that lacked the W3 element (PEmax: 23.8±0.9%; PE3bmax: 38.2±2.6%), and with the terminators that contained the W3 element but used either the synthetic polyA (PEmax: 28.0±3.3%; PE3bmax: 27.0±7.3%;) or SV40 polyA (PEmax: 21.2±6.8%; PE3bmax: 23.8±5.6%; extended data fig. 5e). These three C-terminal PE vector designs were therefore selected for further testing with the N-terminal PE vector in *in vivo* prime editing experiments.

### *In vivo* prime editing at the *Adrb1* and *Dnmt1* locus in the brain

To deliver PE constructs into the brain of newborn C57BL/6 mice, we packaged the selected N-and C-terminal PE expression vectors into AAV capsids of the serotype PHP.eB (Fig 2a,b). In addition to targeting the *Adrb1* locus with epegRNA1 in the PE3b approach, we targeted the *Dnmt1* locus using a pegRNA that has previously been employed in the liver^5^, further stabilized with the tevopreQ_1_ structural motif (extended data table 1). Efficient packaging of the N-and C-terminal PE expression vectors into AAVs was first confirmed by negative staining electron microscopy (Extended data fig. 6a-d), and particles were delivered to the ventricles of neonatal mice via intracerebroventricular (ICV) injection at a dose of 5×10^10^ vg per construct and animal (Fig. 2b). Brains were isolated and cortices were manually dissected at different timepoints post-injection to quantify editing outcomes by deep amplicon sequencing (Fig. 2c). At 5 weeks, mouse cortices showed on average 13.3% editing at the *Adrb1* locus (for W3-synth) and 27.7% editing at the *Dnmt1* locus (for noW3-bGH), without significant differences between the three different minimal terminators utilized in the C-terminal PE constructs (Fig. 2c). In line with these results, RT-qPCR analysis revealed similar C-terminal PE expression levels from the different constructs (Extended data fig. 5e,f). Editing rates were maintained over time at both loci as assessed by deep amplicon sequencing at 10 weeks (17.6±8.6% for W3-synth at the *Adrb1* locus and 32.7±13.1% for noW3-bGH at the *Dnmt1* locus) and 6 months post-injection (14.7±11.6% for W3-synth at the *Adrb1* locus and 34.8±9.8% for noW3-bGH at the *Dnmt1* locus; fig. 2c). Similarly, the formation of indels, primarily caused by the nickase activity of Cas9 rather than integration of the sgRNA scaffold (Extended data fig. 7), did not increase over time and remained at 2.3±1.8% at the *Adrb1* locus and 1.0±0.1% at the *Dnmt1* locus (cortices at 6 months; fig. 2c).

Next, we analyzed variations in editing rates across different brain regions and cell types. As expected from differences in the transduction efficiency of AAV PHP.eB across various brain regions after ICV injections (Extended data fig. 8)^37^, we observed differences in editing rates that roughly reflected the AAV transduction efficiencies in the corresponding regions (Fig. 2d). The use of the hsyn promoter, moreover, limited PE expression to neurons (Extended data fig. 9). Thus, when we analyzed editing rates in neuron-enriched cell populations, we observed a 4- and 2-fold increase in editing rates for *Adrb1* and *Dnmt1*, respectively (Extended data fig. 10). In line with neuron-specific prime editing, we did not observe editing above background in any of the other tested organs except the spinal cord (Fig. 2e). Of note, exchanging the hsyn promoter with the ubiquitously active Cbh promoter^38^ and replacing the W3-bGH terminator (477 bp) on N-and C-terminal AAV vectors with the synthetic polyA (49 bp) resulted in 25.0±1.9% editing at the *Dnmt1* locus in the brain, despite using half the dose compared to previous experiments (2.5×10^10^ vg per construct and animal; fig. 2f), and in editing in the liver (14.2±6.2%) and the heart (2.5±0.9%; fig. 2f) at 10 weeks post-injection (ICV). These data suggest that efficienct prime editing in the brain neither requires the W3 element nor long terminators when the PE is expressed from strong promoters.

*In vitro* in cell lines epegRNA1-SM^CTT^ enabled high on-target editing with substantially lower indel formation rates compared to epegRNA1 (with-or without the PE3b nicking sgRNA), prompting us to also test this pegRNA *in vivo*. epegRNA1-SM^CTT^ was cloned into the N-terminal PEmax expression vector and co-delivered with the C-terminal PEmax expression vector. However, in contrast to our *in vitro* results in Hepa and N2a cells (Fig. 1c), *in vivo* editing rates were substantially lower than with epegRNA1 in the PE3b approach (2.1±1.3% vs. 13.2%±8.3% at 5 weeks post-injection; Fig. 2c,g), and also co-expression of the PE3b-ngRNA together with epegRNA1-SM^CTT^ did not increase editing rates (1.3±0.5%; fig. 2g). Nevertheless, indel rates were significantly lower with epegRNA1-SM^CTT^ than with epegRNA1 and at the levels of saline-treated controls (Fig. 2g). Thus, while editing of the PAM sequence did not enhance editing efficiency, it led to a significant reduction in indel formation. Consistent with this theory, indel rates at the *Dnmt1* locus, where we converted the second G of the PAM into a C, was also at background levels (Fig. 2c,d).

### Installing the *Adrb1*^A187V^ mutation in newborn mice increases activity and exploration

The *Adrb1*^A187V^ variant has recently been shown to induce increased activity and short sleep in humans and mice^7^. Hence, we next assessed if installing *Adrb1*^A187V^ in the brain of newborn mice changed their activity. First, we monitored mice in their home cage during the light-and dark cycle using infrared motion sensors 4 weeks after injection of epegRNA1 in the PE3b approach. In line with previous reports^7^, treated animals displayed longer active periods, with a significant increase compared to saline-injected controls or *Dnmt1*-targeting AAV controls during the dark cycle (Fig. 3a). Next, we employed the open field (OF) test to assess whether locomotor activity was increased (Fig. 3b). Our results showed that treated mice covered significantly more distance (46.9±9.0m vs. 38.5±9.4m in the light cycle and 50.4±7.9m vs. 44.3±8.2m in the dark cycle; fig. 3c) and moved at higher velocity than control animals (14.6±0.8 cm/s vs. 13.4±1.3cm/s in the light and 16.5±1.2cm/s vs. 14.5±1.5cm/s in the dark cycle; fig. 3c). We further observed an increase in the frequency of wall-supported rearings in treated animals, both in the OF (Fig. 3c) and when animals were placed in an unfamiliar environment (Fig. 3d), indicating enhanced exploratory behavior in treated animals^39^. Notabtly, we did not detect a significant increase in stress-or anxiety-related responses in the OF^40, 41^ or light-dark (LD) transition test ^42^ (Extended data fig. 11). Since β-ARs are also important for memory^14^, we further assessed memory performance using the novel object (NO) recognition test (Extended data fig. 11c). Our data did not indicate signs of memory impairment in response to the *Adrb1*^A187V^ mutation (Extended data fig. 11c). In fact, although not significant, Adrb1-treated mice spent on average slightly more time with the novel object (58.0±8.7% vs. 51.8±6.7%; extended data fig. 11d), suggesting that the *Adrb1*^A187V^ variant might have a beneficial impact on learning and memory.

Both on-target editing (9.1±4.1%) and indel formation (2.0±0.9%) was detected in cortices of treated animals at experimental endpoints (Extended data fig. 12a). Since *Adrb1*^A187V^ is a dominant negative allele that leads to reduced protein stability^7^, indel mutations may also contribute to the observed behavioral phenotype. To exclude that the effects are solely a consequence of indel formation, we repeated the OF test with newborn animals treated with PEmax and epegRNA-SM^CTT^, which did not show indels above background (Extended data fig. 12a). In line with our previous observations (Fig. 3c), also epegRNA-SM^CTT^-treated mice displayed a significant increase in locomotor activity during the light phase (43.9±6.0m vs. 31.2±5.7m; extended data fig. 12b), with a direct correlation between the traveled distance and *Adrb1*^A187V^ editing rates but not indel rates (Extended data fig. 12c). Next, we assessed if the observed phenotype could be related to pegRNA-depedent off-targets effects. We used CHANGE-seq^43^ to experimentally identify potential off-target binding sites of the Adrb1 protospacer, which were then analyzed by deep sequencing. Importantly, in the top 5 identified off-target sites we observed no differences in indels or substitutions between PE-and saline-treated animals (Extended data fig. 13). Taken together, our data indicate that installation of the *Adrb1*^A187V^ mutation in newborn mice induced a behavioral phenotype.

### Installing *Adrb1*^A187V^ in the mPFC of adult mice increases general activity and exploration

Next we analyzed which brain regions might be associated with the alterations in animal behavior, and separately quantified editing rates in regions where *Adrb1* is expressed^7, 44–46^ (Extended data fig. 14a). Deep amplicon sequencing revealed that editing rates were several fold higher in the medial prefrontal cortex (mPFC) (12.2±4.5%) compared to the other areas with *Adrb1* expression, including the hippocampal formation (dorsal: 2.7±1.7%, ventral: 3.1±1.7%), lateral septal nucleus (1.8±1.0%), caudate putamen (3.6±1.4%), amygdala (1.3±1.2%), thalamus (3.0±0.9%), paraventricular nucleus (0.6±0.3%), midbrain (1.5±1.4%), dorsal pons (0.4±0.2%), and medulla oblongata (0.4±0.4%) (Extended data fig. 14b). These results prompted us to assess if installing *Adrb1*^A187V^ directly in the mPFC via intracranial AAV injection in adult mice could lead to similar phenotypes (Fig. 4a). Deep amplicon sequencing of tissue isolated from the mPFC of mice treated with PEmax and epegRNA-SM^CTT^ revealed editing rates of 5.8±2.6% (Fig. 4b). This value, however, is likely an underestimation of editing in mPFC neurons, becaused dissected tissues also contained surrounding regions of the mPFC and glial cells. 4 weeks post-injection we performed behavioral tests with treated and untreated control animals. While, in contrast to newborns treated via ICV injection, locomotor activity and velocity were comparable between treated and control animals (Fig. 4d), the active time in the home cage as well as the frequency of wall-supported rearings was significantly increased (Fig. 4c,d). Moreover, similar to our observations in newborn animals, we did not observe an increase in anxiety-related behavior in the OF, but a trend for enhanced memory performance in the NO recognition test (Fig. 4f). Together, these results indicate that the behavioral phenotypes observed in ICV-treated newborns are partly caused by *Adrb1*^A187V^ positive neurons of the mPFC, although it is likely that editing in other brain areas with *Adrb1* expression also contributed to the observed behavioral phenotype.

## Discussion

In our study we developed an approach for *in vivo* prime editing in the brain. Delivery of optimized AAV vectors encoding for intein-split PEs into the brain of newborn mice resulted in editing rates of up to 44.0% at the *Dnmt1* locus and 28.1% at the *Adrb1* locus in the cortex. Introducing the *Adrb1*^A187V^ mutation was greatly enhanced using the PE3b approach, in which a second ngRNA is employed to cut the unedited strand after editing at the PAM strand occurred. In addition, we found that prime editing precision can be increased by introducing a silent mutation in the PAM, which circumvents retargeting of the edited locus by the PE complex. Notably, at the *Adrb1*^A187V^ site the integration of a silent mutation in the PAM led to decreased editing rates *in vivo*. However, it is uncertain whether this observation can be generalized, as a comprehensive evaluation across multiple sites would be required.

In line with previous studies analyzing the behavior of heterozygous and homozygous *Adrb1*^A187V^ mice^7^, introducing the *Adrb1*^A187V^ mutation into newborn mice via prime editing led to increased activity, locomotion, and exploratory behavior, with a direct correlation between editing rates and active time in the home cage or traveled distance in the OF (Extended data fig. 12; extended data fig. 15). Increased activity has also been observed in animals where *Adrb1*^A187V^ was specifically introduced into neurons of the mPFC, suggesting that this region plays an important role for *Adrb1*^A187V^ effects on behavior.

Together our experiments provide proof-of-concept for prime editing in the brain to modulate neural circuits and animal behavior. Since β1-ARs are important for various brain functions, the developed tools could prove valuable for modulating β1-AR function in specific brain regions in the context of health and disease. Moreover, *in vivo* prime editing in the brain could also be employed to correct various genetic brain disorders, including psychiatric disorders or neurodegenerative diseases.

## Methods

### Generation of plasmids

To generate pegRNA plasmids, annealed spacer, scaffold, and 3’ extension oligos were cloned into the BsaI-digested pU6-pegRNA-GG- (Addgene #132777), pU6-tevopreQ1-GG- (Addgene #174038) or pU6-tmpknot-GG-acceptor plasmid (Addgene #174039) using Golden Gate assembly as previously described^1, 11^. ngRNA and shRNA plasmids were generated by ligating annealed and phosphorylated oligos into a BsmBI-digested lentiGuide-Puro (Addgene #52963) or an EcoRI-digested pLKO.1 backbone using T4 DNA ligase (Addgene #8453). To generate intein-split PE plasmids, inserts were either ordered as gBlocks from Integrated DNA Technologies (IDT) or amplified from pCMV-PE2 (Addgene #132775) or pCMV-PEmax (Addgene #174820) plasmids using PCR. Inserts were cloned into the NotI-and EcoRI-digested pCMV-PE2 backbone using HiFi DNA Assembly Master Mix [New England Biolabs (NEB)]. To generate PiggyBac reporter plasmids for the *Adrb1* locus, inserts with homology overhangs for cloning were ordered from IDT and cloned into the XbaI-and EcoRI-digested pPB-Zeocin backbone using HiFi DNA Assembly Master Mix (NEB). To prepare plasmids for AAV production, inserts with homology overhangs were either ordered as gBlocks (IDT) or generated by PCR. Inserts were cloned into XbaI-and NotI-digested AAV backbones using HiFi DNA Assembly Master Mix (NEB). All PCRs were performed using Q5 High-Fidelity DNA Polymerase (NEB). The identity of all plasmids was confirmed by Sanger Sequencing. Primers used for cloning all plasmids are listed in extended data tables 1-3. Amino acid sequences of intein-split PEmax p.713/p.714 constructs are listed in extended data table 4.

### Cell culture transfection and genomic DNA preparation

HEK293T [American Type Culture Collection (ATCC) CRL-3216] and Hepa1-6 (ATCC CRL-1830) cells were maintained in Dulbecco’s modified Eagle’s medium (DMEM) plus GlutaMAX (Thermo Fisher Scientific), supplemented with 10% (v/v) fetal bovine serum (FBS) and 1% penicillin/streptomycin (Thermo Fisher Scientific) at 37°C and 5% CO_2_. Neuro2a (ATCC CCL-131) cells were maintained in Eagle’s Minimum Essential Medium (EMEM), supplemented with 10% (v/v) FBS and 1% penicillin/streptomycin. Cells were passaged every 3 to 4 days and maintained at confluency below 90%.

Cells were seeded in 96-well cell culture plates (Greiner) and transfected at 70% confluency using 0.5 μl Lipofectamine^TM^ 2000 (Thermo Fisher Scientific). If not stated otherwise, 300 ng of PE, 100 ng of pegRNA, 40 ng of nicking sgRNA (where indicated), and 150 ng of dnMLH1 (Addgene #174824) were used for transfection. When intein-split PEs were transfected, 300 ng of each PE half was used. Cells were incubated for 3 days after transfection.

Genomic DNA from cells was isolated using a lysis buffer (10 mM Tris-HCl, 2 % Triton™ X-100, 1 mM EDTA, proteinase K [20 mg/mL]; Thermo Fisher Scientific). Cells were lysed at 60°C for 1 h, followed by 10 min at 95°C. Lysates were further diluted with 60 µL of dH_2_O. Genomic DNA from mouse tissues was isolated by phenol/chloroform extraction. First, tissue samples were incubated overnight in lysis buffer (50 mM Tris-HCl pH 8.0, 100 mM EDTA, 100 mM sodium chloride, and 1% SDS; Thermo Fisher Scientific) at 55°C and 300 rpm. Subsequently, phenol/chloroform/isoamyl alcohol (25:24:1, Thermo Fisher Scientific) was added and samples were centrifuged (5 min, maximum speed). The upper phase was transferred to a clean tube and DNA was precipitated using 100% ethanol (Sigma-Aldrich). Samples were centrifuged (5 min, maximum speed) and pellets were washed using cold 70% ethanol (−20°C). Washed pellets were dried at 55°C for 10 min and resuspended in 100 µL of dH2O.

### Generation of the reporter and MMR-deficient cell lines

To generate Adrb1 reporter cell lines with the PiggyBac transposon, HEK, Hepa, and N2a cells were seeded into a 48-well cell culture plate (Greiner) and transfected at 70% confluency with 225 ng of the PiggyBac-transposon and 25 ng of the transposase using Lipofectamine 2000 (Thermo Fisher Scientific) according to the manufacturer’s instructions. Three days after transfection, cells were enriched for 10 days using Zeocin selection [150 μg/ml].

MMR-deficient Adrb1 reporter cell lines were generated using lentivirus transduction of shRNAs, targeting murine or human *Mlh1/MLH1* or *Msh2/MSH2*. For lentivirus production, HEK293T cells were seeded into a 6-well cell culture plate (Greiner) and transfected with 1500 ng of cargo, 400 ng of VSV-G (Addgene #8454), and 1100 ng of PAX2 (Addgene #12260) plasmid using polyethylenimine (PEI, Polysciences). The cell culture medium was exchanged 12 h after transfection and the virus was harvested 24 h later. Supernatants containing lentiviral particles were added to HEK, Hepa, and N2a Adrb1 reporter cell lines, which were seeded into 6-well cell culture plates (Greiner) one day prior to transduction. Transduced cells were enriched for 7 days with Puromycin [2.5 µg/mL].

### AAV production

For the production of a pseudo-typed vector (AAV2 serotype PHP.eB) expressing EGFP under the control of the Cbh promoter, 2×10^8^ HEK293T cells were seeded per 150 mm dish (5 dishes in total) 24 h prior to transfection. For transfection, the helper plasmid (25 μg per dish), capsid plasmid (15 μg per dish) and cargo plasmid (9 μg per dish) were mixed with serum-free DMEM (Thermo Fisher Scientific) and polyethylenimine (PEI, Polyscience) was added in a 1:3 ratio (1 mg of DNA to 3 mg of PEI). The mix was incubated for 20 min at RT and then added to the cells. After 5 days of incubation at 37°C and 5% CO2, cells were harvested and centrifuged for 15 min at 1’500×g in a conical corning flask (Thermo Fisher Scientific). 150 mL of the supernatant were mixed with 22 mL of NaCl (5M) and 30 mL of PEG8000 (VWR) in a new corning flask. The cell pellet was resuspended in 4 mL of resuspension buffer (150 mM NaCl, 50 mM Tris-HCl, pH 8.0) and homogenized using a Precellys Evolution homogenizer (2 cycles: 5’000 rpm for 45 sec with 15 sec break). 300 units of benzonase (Sigma-Aldrich) were added to the disrupted cells and the mixture was incubated for 30 min at 37°C in a water bath. After centrifugation at 5’000×g for 1h, the supernatant was collected and mixed with the supernatant after harvesting. AAV particles in this mixture were precipitated for 2 days at 4°C, followed by centrifugation at 10’000×g for 30 min. The supernatant was discarded and the AAV particles were washed using 4 mL of resuspension buffer. Particles were resuspended in 1.5 mL of NaCl [5M]. Next, four fractions of OptiPrep GradientDensity medium (Sigma-Aldrich) were prepared (15%, 25%, 40%, and 60%). The most concentrated fraction was prepared at the bottom of the ultracentrifugation tube and least concentrated fraction at the top of the tube. The virus suspension was added at the top of the tube and the gradient was ultracentrifuged at 65’000 rpm at 15°C for 2 h. AAV particles were harvested from the 40% gradient fraction and filtered using a pre-washed 100 kDa Amicon (Vivaspin). Virus particles were subsequently washed multiple times with PBS (pH 7.4, Thermo Fisher Scientific) and physical titers were measured using a Qubit 3.0 fluorometer and the Qubit dsDNA HS assay kit (Thermo Fisher Scientific).

All other pseudo-typed vectors (AAV2 serotype PHP.eB) were produced by the Viral Vector Facility of the Neuroscience Center Zurich. Briefly, AAV vectors were ultracentrifuged and diafiltered. Physical titers (vector genomes per milliliter, vg/mL) were determined using a Qubit 3.0 fluorometer (Thermo Fisher Scientific) as previously published^47^. The identity of the packaged genomes of each AAV vector was confirmed by Sanger sequencing.

### Negative staining and electron microscopy

First, carbon-coated electron microscopy (EM) grids (200 mesh, Quantifoil) were glow-discharged. Grids were briefly washed with a drop of 0.01% bovine serum albumin (BSA, Sigma-Aldrich). Subsequently, 2 μL of the sample was applied to the grid and incubated for 5 min. Excess liquid was removed from the edge of the grid with filter paper (Whatman). Next, grids were washed with 1 mM EDTA (Sigma-Aldrich), followed by staining with 0.5% uranyl acetate for 1 min. The liquid was again removed from the edge of the grid with filter paper and grids were dried overnight before imaging. Data were collected using an FEI Talos 120 kV transmission electron microscope (Thermo Fisher Scientific) equipped with a digital CMOS camera. Micrographs of several grid squares were collected for each AAV preparation to determine the ratio of fully packaged, partially packaged, and empty AAV particles. Data were quantified using MAPS (Thermo Fisher Scientific) and Fiji^48^.

### Animal studies

Animal experiments were performed in accordance with protocols approved by the Kantonales Veterinäramt Zürich and in compliance with all relevant ethical regulations. C57BL/6J mice were housed in a pathogen-free animal facility at the Institute of Pharmacology and Toxicology of the University of Zurich. Mice were kept in a temperature-and humidity-controlled room on a 12-hour light-dark cycle. Mice were fed a standard laboratory chow (Kliba Nafag no. 3437 with 18.5% crude protein).

Unless stated otherwise, newborn mice (P1) received 5.0 × 10^10^ vg per animal and construct via intracerebroventricular injection (ICV). Adult male C57BL/6J mice at P50-P60 were used to perform surgeries for the delivery of PE-AAVs (dose of 3.0 × 10^8^ vg per hemisphere). Buprenorphine [0.1 mg/kg body weight] was administered to mice subcutaneously 30 min prior to surgery. Animals were anesthetized using isoflurane (5% isoflurane with 1000 mL/min in 100% O_2_) and placed into a stereotaxic mouse frame on a warming surface to maintain body temperature. Anesthesia was maintained at 1.5-2.5% isoflurane with 400 mL/min in 100% O_2_ during surgeries. AAVs were injected bilaterally into the medial prefrontal cortex (mPFC) at the coordinates relative to bregma: 1.8 mm anteroposterior (AP); ± 0.4 mm mediolateral (ML); −1.8 mm dorsoventral (DV) and dorsal pons (dPons, −5.1 mm AP; ± 0.5 mm ML; −3.5 mm DV). 400 nL injections were performed using a glass needle at a speed of 50 nL/min. The needle was slowly removed 3 min after injection and the wound was sutured using Vicryl 5-0 suture (Ethicon).

Behavior experiments and activity monitoring were performed at 4 weeks post injection. Newborn mice were euthanized at 5, 10 or 24 weeks of age. Adult mice were euthanized at 16 weeks of age.

### Behavioral assays

For the open field tests, a 50×50×50 cm chamber made of black plastic walls and a white floor was used. Mice were placed in the center of the open field and their activity was automatically recorded for 10 min from the top (C270 HD Webcam, Logitech). Tests were conducted under background dim illumination (intensity 30 Lx) in the light phase, and under dim red light (intensity 30 Lx) in the dark phase. All tests were performed at least 2 hours after the respective light phase onset and finished at least 2 h before the onset of the next light/dark cycle. To minimize the stress of mice, animals were brought into the experimental room in their home cages at least 1 h prior to the test. Locomotor activity was automatically quantified as distance traveled during a 10 minutes period using a custom-written MATLAB script. The average speed of a mouse was calculated as the distance covered during the running time divided by the time the mouse spent running. Running ‘episodes’ were automatically detected as the time intervals when the instant velocity of the mouse was higher than a given threshold of 5 cm/s. Instant velocity of the mouse was calculated for a sliding window of 4 frames.

For the assessment of anxiety-like behaviors, time spent in the center of the OF arena was automatically quantified using a custom-written MATLAB script or time spent in the light compartment in the light-dark transition test (20×20×20 cm each) and transitions from the dark to the light compartment were manually quantified. The dark box was covered with an opaque lid and the light box was covered with a transparent top during the 10 min experiment.

For the assessment of explorative behavior, wall-supported rearing in the OF was manually quantified in the light and dark phases. For additional analysis of explorative behavior, mice were placed in a cylinder and recorded for 10 min from the side. Wall-supported rearing in the cylinder was manually quantified in the light phase.

For analysis of memory performance, the same OF arena as described above was used. On day 1, animals were habituated to the arena for 10 min. On day 2, animals were placed into the arena with two identical for 10 min. On day 3, one of the objects was replaced and the number of approaches of the novel object was manually quantified for 10 min.

For monitoring the activity of the mice using motion sensors, animals were single-housed and an infrared sensor was placed on top of the home cage. After 1-week of habituation, activity was recorded for at least 5 consecutive days and data were analyzed using ClockLab (Actimetrics).

### Trans-cardiac perfusion, brain isolation, and dissection of brain regions

Sodium Pentobarbitol (Kantonsapotheke Zürich) was injected via intraperitoneal injection at a dose of 100 mg/kg. Complete anesthesia was confirmed by the absence of a toe pinch reflex. Mice were placed on the perfusion stage inside a collection pan and the peritoneal cavity was exposed. The diaphragm was cut through laterally and the rib cage was cut parallel to the lungs, creating a chest “flap”. The flap was clamped in place using a hemostat (Fine Science Tools) and a 25 gauge needle (Sterican), attached to silicon tubing and a peristaltic pump, was inserted into the left ventricle. The right atrium was cut for drainage. Animals were first perfused with ice-cold PBS (Thermo Fisher Scientific) at a rate of 10 mL/min, followed by perfusion with ice-cold fixative (4% paraformaldehyde, PFA, Sigma-Aldrich). When brains were used for a single cell, DNA, RNA, or protein isolation, perfusion was performed exclusively with PBS. Once the perfusion was complete, mice were decapitated and the skull was removed with scissors and tweezers without inflicting damage to the underlying tissue. The brain was removed using a spatula.

For histology, PFA-perfused brains were post-fixated in 4% PFA for 4h, followed by overnight incubation in 30% sucrose. For the dissection of brain regions, PBS-perfused brains were first rinsed in PBS and then cut into 1 mm slices using an acrylic mouse brain matrix (AgnThos) and razor blades. The olfactory bulb, cortex, hippocampus, striatum, thalamus, hypothalamus, midbrain, hindbrain, and cerebellum were identified based on the mouse brain atlas^49^. To dissect *Adrb1*-expressing regions with high precision, 60 μm sections of PBS-perfused brains were prepared, and the region of interest was isolated under a stereomicroscope using the mouse brain atlas^49^.

### RNA isolation and RT-qPCR

RNA was isolated from cultured or isolated cells or snap-frozen brain tissues using the RNeasy Mini Kit (Qiagen) or the RNeasy Lipid Tissue Mini Kit (Qiagen) according to the manufacturer’s instructions. RNA (1000 ng input) was subsequently reverse-transcribed to cDNA using random primers and the GoScript RT kit (Promega). RT-qPCR was performed using FIREPoly qPCR Master Mix (Solis BioDyne) and analyzed using a Lightcycler 480 system (Roche). Fold changes were calculated using the ΔCt method. Primers used for RT-qPCR are listed in extended data table 5.

### Protein isolation and western blot

Protein was isolated from cultured cells using radioimmunoprecipitation (RIPA) assay buffer (150 mM Tris pH 8.0, 150 mM NaCl, 0.1% SDS, 0.5% sodium deoxycholate, 1% NP-40; Thermo Fisher Scientific), supplemented with protease inhibitor cocktail (Roche). Protein concentrations of all samples were determined using the Pierce Bicinchoninic Acid (BCA) Protein Assay Kit (Thermo Fisher Scientific).

Equal amounts of protein (*in vitro* samples: 30 μg) were separated by SDS-polyacrylamide gel electrophoresis (Thermo Fisher Scientific) and transferred to a 0.45-μm nitrocellulose membrane (Amersham). Membranes were incubated with rabbit anti-Adrb1 (1:1,000; cat. no. ab85037, Abcam) and mouse anti-actin beta (1:2,000; cat. no. ab8226; Abcam). Signals were detected by fluorescence using IRDye-conjugated secondary antibodies (LI-COR Biosciences) and a LICOR Odyssey ® DLx imaging system. All antibodies are listed in extended data table 6.

### Single-cell isolation by MACS

PBS-perfused brains were cut into small pieces and dissociated using the Adult Brain Dissociation Kit and gentleMACS Octo Dissociator with heaters (Miltenyi Biotec) according to the manufacturer’s instructions. Cell debris and myelin was subsequently removed using a debris removal solution (Miltenyi Biotec). The single-cell suspension was used for the isolation of neurons using the Adult Neuronal Isolation kit (Miltenyi Biotec) according to the manufacturer’s instructions.

The identity of all fractions was confirmed by flow cytometry and RT-qPCR. Briefly, cells were resuspended in FACS buffer (PBS supplemented with 2% FBS and 2 mM EDTA). Fc blocking reagent (1:50 in FACS buffer) was added to all samples and incubated on ice for 20 min. Samples were subsequently labeled with primary antibodies (ACSA-2, O4, CD11b, Biotin, eFluor789, in FACS buffer) for 1 h at 4°C (in the dark). Cell suspensions were washed three times in FACS buffer and filtered through a 35-μm nylon mesh cell strainer snap caps (Corning) and kept on ice until analysis. For each sample, 10,000-50,000 events were counted on an LSRFortessa (BD Biosciences) using the FACSDiva software version 8.0.1 (BD Biosciences). Experiments were performed with three replicates/mice. The gating strategy is shown in extended data fig. 8. RT-qPCR primers and antibodies are listed in extended data tables 4 and 5.

### Amplification for deep sequencing

Genomic DNA from cultured cells or brain tissues was isolated by direct lysis (cells) or phenol/chloroform extraction (brain tissue). *Adrb1-* or *Dnmt1*-specific primers were used to generate targeted amplicons for deep sequencing. Input genomic DNA was first amplified in a 10μL reaction for 30 cycles using NEBNext High-Fidelity 2×PCR Master Mix (NEB). Amplicons were purified using AMPure XP beads (Beckman Coulter) and subsequently amplified for eight cycles using primers with sequencing adapters. Approximately equal amounts of PCR products were pooled, gel purified, and quantified using a Qubit 3.0 fluorometer and the dsDNA HS Assay Kit (Thermo Fisher Scientific). Paired-end sequencing of purified libraries was performed on an Illumina Miseq. Primers for deep sequencing are listed in extended data table 7.

### HTS data analysis

Sequencing reads were first demultiplexed using the Miseq Reporter (Illumina). Next, amplicon sequences were aligned to their reference sequences using CRISPResso2^50^. Prime editing efficiencies were calculated as percentage of (number of reads containing only the desired edit)/(number of total aligned reads). Indel rates were calculated as percentage of (number of indel-containing reads)/(total aligned reads). Reference sequences are listed in extended data table 8.

### Immunohistochemistry

PFA-fixed brain tissues were frozen on dry ice and cut into 40 μm-thick sections using a microtome. Sections were blocked in PBS supplemented with 2% normal donkey serum (cat. no. ab7475, abcam) and 0.3% Triton X-100 (Sigma-Aldrich) for 1 h. Brain sections were incubated with primary antibodies overnight at 4°C (mouse-NeuN, 1:500, abcam ab177487; rabbit-Cas9, 1:1,000, Cell Signaling clone D8Y4K; chicken-GFAP, 1:1’500, abcam ab95231). Donkey anti-mouse-568 (1:500), donkey anti-chicken-647 (1:500) and donkey anti-rabbit-488 (1:1,000; all from Jackson ImmunoResearch) were used as secondary antibodies and sections were counterstained with 4′,6-diamidino-2-phenylindole (DAPI, Sigma-Aldrich). Mounting was performed using Prolong Gold Antifade Mountant (Thermo Fisher Scientific). Confocal images were taken with a Zeiss LSM 800 or a Zeiss AxioScan.Z1 slidescanner and analyzed with Fiji^48^. Antibodies are listed in extended data table 6.

### Statistical analysis

All statistical analyses were performed using GraphPad Prism 9.0.0 for macOS. If not stated otherwise, data are represented as biological replicates and are depicted as means±s.d. Statistical analyses are always indicated in the corresponding figure legends. Likewise, sample sizes and the statistical tests performed are described in the respective figure legends. The data were tested for normality using the Shapiro-Wilk test if not stated otherwise. Unpaired two-tailed Student’s *t*-tests were performed followed by the appropriate post hoc test when more than two groups were compared. For all analyses, *p*<0.05 was considered statistically significant.

## Acknowledgements

We thank the Functional Genomics Center Zurich for technical support and access to instruments at the University of Zurich. Furthermore, we acknowledge Andres Käch, José María Mateos Melero, and Johannes Riemann from the Center for Microscopy and Image Analysis at the University of Zurich for support with negative staining and TEM imaging of AAV preparations. We thank Steven Brown for his constructive feedback in the course of this study and critical assessment of the data. We thank Martha Gjikolai and Sara Pierre-Ferrer for their help with infrared activity monitoring. We thank Hanns Ulrich Zeilhofer for sharing lab equipment and access to experimental rooms for behavior analyses. Members of the Schwank, Patriarchi, and Brown labs are acknowledged for discussions and comments on the manuscript. This work was supported by the Swiss National Science Foundation (SNSF) grant no. 310030_185293 (to G.S.), Novartis Foundation for Medical-Biological Research no. FN20- 0000000203 (to D.B.), SNSF Spark fellowship no. 196287 (to D.B.), the URPP Itinerare (to G.S. and to D.B.) and the Helmut Horten Foundation (to G.S.).

## Author contributions

D.B. and G.S. designed the study. D.B., L.T., Y.W., E.I., T.R., L.S., and S.J. performed and analyzed *in vitro* experiments. E.I. produced and quantified AAV-PHP.eB pCbh-GFP particles used for quantification of transduction efficiencies across the brain. M.W., D.B., and L.T. performed and analyzed *in vivo* experiments. J.M. sectioned, stained, and imaged AAV-transduced brains. D.B. and G.S. wrote the manuscript with input from all coauthors. All authors reviewed the manuscript.

## Data availability

All data associated with this study are present in the paper. Illumina sequencing data will be made available at the Gene Expression Omnibus (GEO) database upon publication.

## Supplementary information

The supplementary information file contains extended data figures 1-15 and extended data tables 1-8.

## Correspondence

Correspondence should be addressed to G. Schwank.

