## Supplementary materials for "Prime editing of the β_1_ adrenoceptor in the brain reprograms mouse behavior"

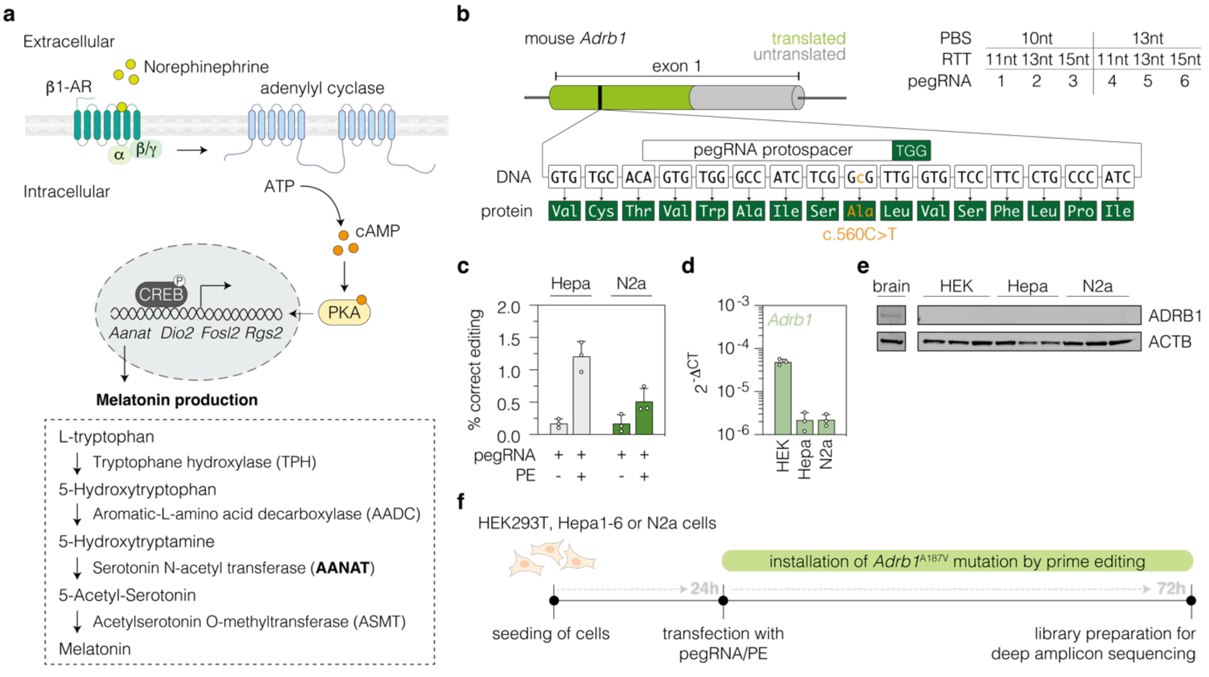


**Extended data figure 1: Establishment of Adrb1 reporter cell lines.** (**a**) Schematic representation of β1-AR-mediated signaling and its potential link to wakefulness and sleep (created based on[51]). (**b**) Depiction of the *Adrb1* gene and pegRNA designs used at the target site (c.560C>T; p.A187V). (**c**) Prime editing rates of pegRNA1 at the endogenous *Adrb1* locus in murine Hepa and N2a cell lines. (**d,e**) *Adrb1* transcript (d) and protein levels (e) in HEK, Hepa, and N2a cells. *Adrb1* protein levels in brain lysate are shown as a positive control. (**f**) Schematic representation of the experimental timeline of prime editing experiments performed in this study. β1-AR, $\beta_1$ adrenoceptor; ATP, adenosine triphosphate; cAMP, cyclic adenosine monophosphate; PKA, protein kinase A; CREB, cAMP responsive element binding protein; LTD/RTD, left/right transposable domains; EF1α, elongation factor alpha; bGH, bovine growth hormone; SV40, Simian virus 40; HSV-tk, Herpes simplex virus thymidine kinase; bp, base pairs. Data are represented as means±s.d. of three independent experiments (b) or three passages of cells (c,d).

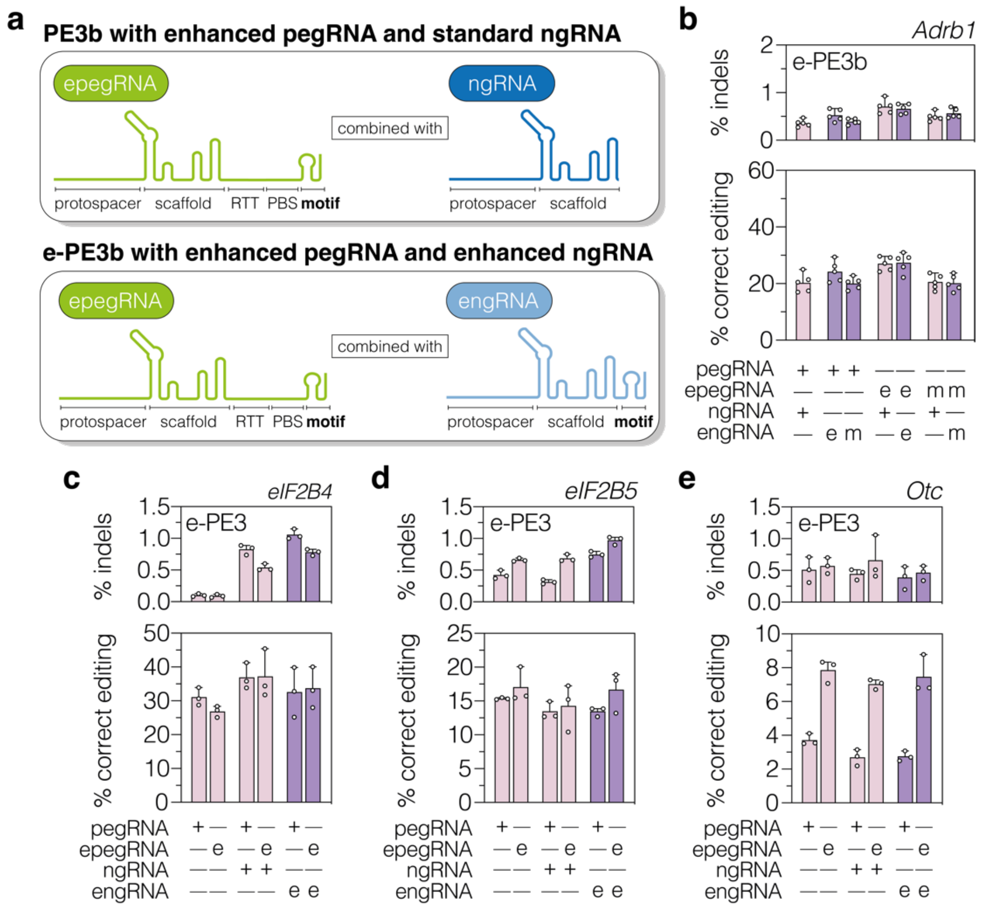


**Extended data figure 2: Enhanced ngRNAs display similar *in vitro* performance to regular ngRNAs.** (**a**) Schematic representation of the PE3b (epegRNA and ngRNA) and e-PE3b approach (epegRNA and engRNA). (**b-e**) e-PE3b editing and indel rates at the *Adrb1* (b), *eIF2B4* (c), *eIF2B5* (d), *and Otc* (e) locus. Data are represented as means±s.d. of at least three independent experiments and were analyzed using a two-tailed Student's t-test with Welch's correction. If not indicated, differences were not statistically significant (*P>0.05*). e, tevopreQ$_1$ motif; m, tmpknot motif.

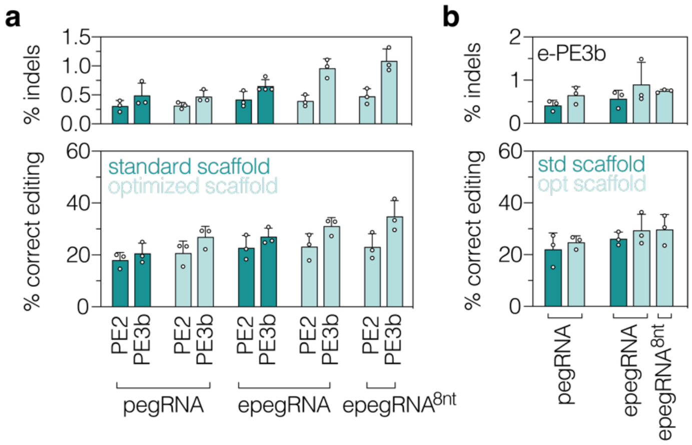


**Extended data figure 3: Effect of scaffold and epegRNA optimizations on *in vitro* prime editing efficiencies.** (a,**b**) Comparison of *in vitro* editing and indel rates of pegRNA1, epegRNA1, and epegRNA$^{8nt}$ (8nt-long linker between the PBS and the tevopreQ$_1$ pseudoknot) with the standard (dark green) or the optimized scaffold (light green). The non-modified PE3b ngRNA was used in (a) and the enhanced PE3b ngRNA was used in (b). Data are represented as means±s.d. of three independent experiments and were analyzed using a two-tailed Student's t-test with Welch's correction. If not indicated, differences were not statistically significant (*P>0.05*).

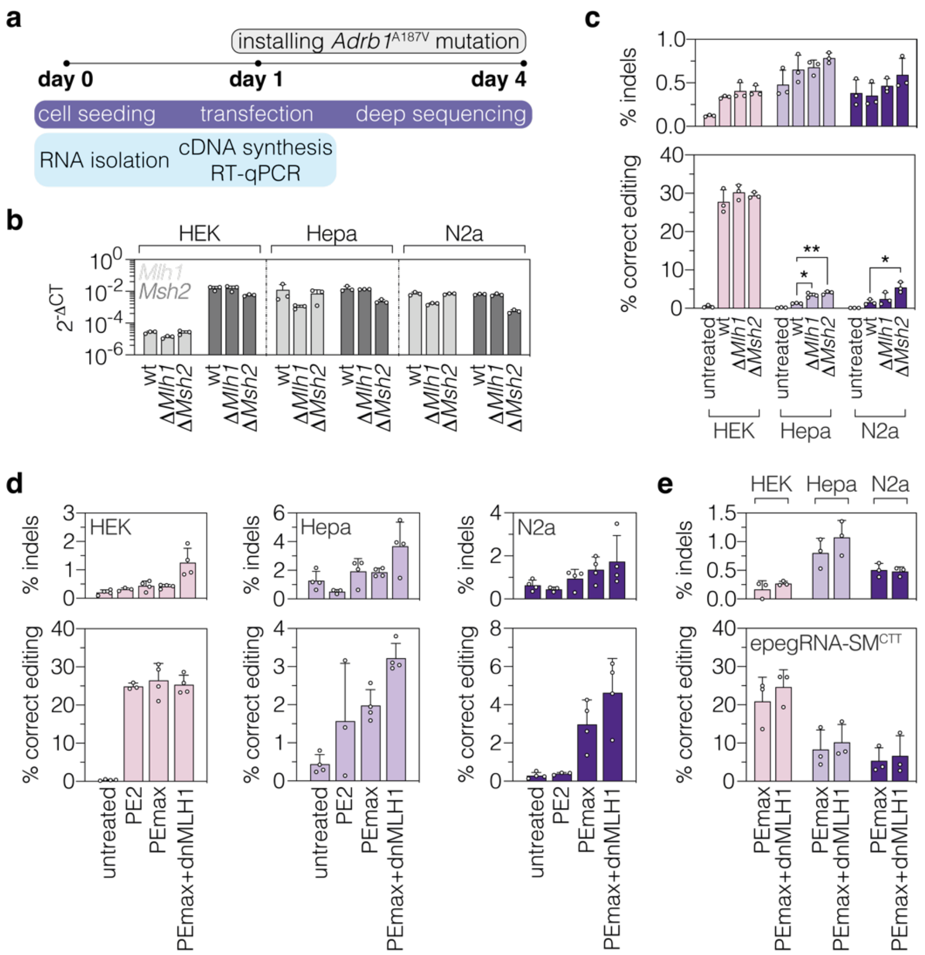


**Extended data figure 4: MMR evasion enhance *in vitro* prime editing rates in MMR proficient cell lines.** (**a**) Schematic representation of the experimental workflow of editing experiments and parallel validation of *Mlh1* and *Msh2* downregulation by RT-qPCR. (**b**) *Mlh1* and *Msh2* transcript levels in partially MMR deficient HEK and MMR proficient Hepa and N2a Adrb1 reporter cells (labelled as wild-type, wt) and Δ*Mlh1*/Δ*Msh2* Adrb1 reporter cell lines (labelled as Δ*Mlh1*/Δ*Msh2*). Transcript levels were normalized to *Gapdh*. (**c**) Editing and indel rates of epegRNA1 and PE2 at the *Adrb1* locus upon treatment with Lentivirus-integrated shRNAs targeting *Mlh1* or *Msh2*. (**d**) Comparative editing and indel rates of epegRNA1 complexed with the original PE or the optimized PEmax variant upon co-transfection of dnMLH1 or a PE3b-ngRNA in HEK, Hepa, and N2a Adrb1 reporter cells. (**e**) Editing and indel rates of epegRNA-SM$^{CTT}$ with co-expression of dnMLH1. Data are displayed as means±s.d. of at least three independent experiments and were analyzed using a two-tailed Student's t-test with Welch's correction (**P<0.05*; ***P<0.005; ***P<0.0005; ****P<0.0001*). If not indicated, differences were not statistically significant (*P>0.05*).

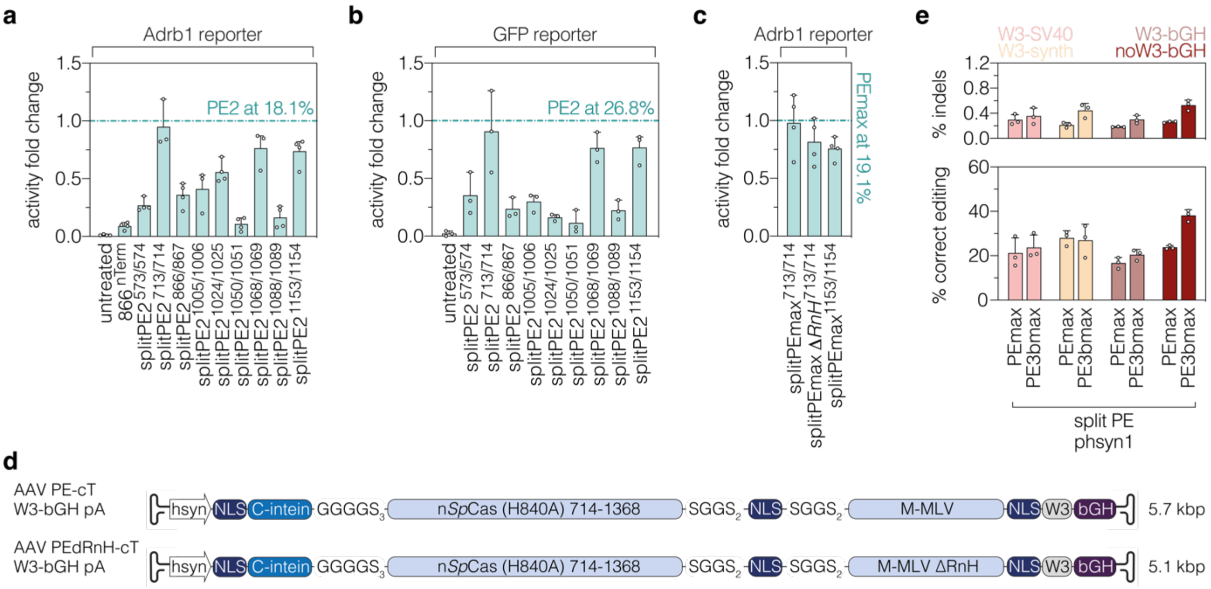


**Extended data figure 5: *In vitro* refinement of intein-split PE variants and AAV designs for the murine brain.** (**a-c**) Relative performance of intein-split PE variants in HEK Adrb1 (a,c) and GFP (b) reporter cell lines. Performance was normalized to full-length PE2 or PEmax (indicated as a dashed line). (**d**) Schematic representation of oversized c-terminal (cT) PE AAV constructs and the corresponding lengths in kilobase pairs (kbp, including ITRs). Constructs are not depicted to scale. (**e**) Comparison of editing and indel rates of intein-split PEmax/PE3bmax AAV constructs under the control of the hsyn promotor. Data are represented as means±s.d. of at least three independent experiments. PEdRnH, PE lacking delta RnaseH domain; phsyn, human synapsin promoter; NLS, nuclear localization signal; W3, woodchuck hepatitis virus post-transcriptional regulatory element; SV40, Simian virus 40; synth, synthetic polyA, bGH, bovine growth hormone; RT, reverse transcriptase.

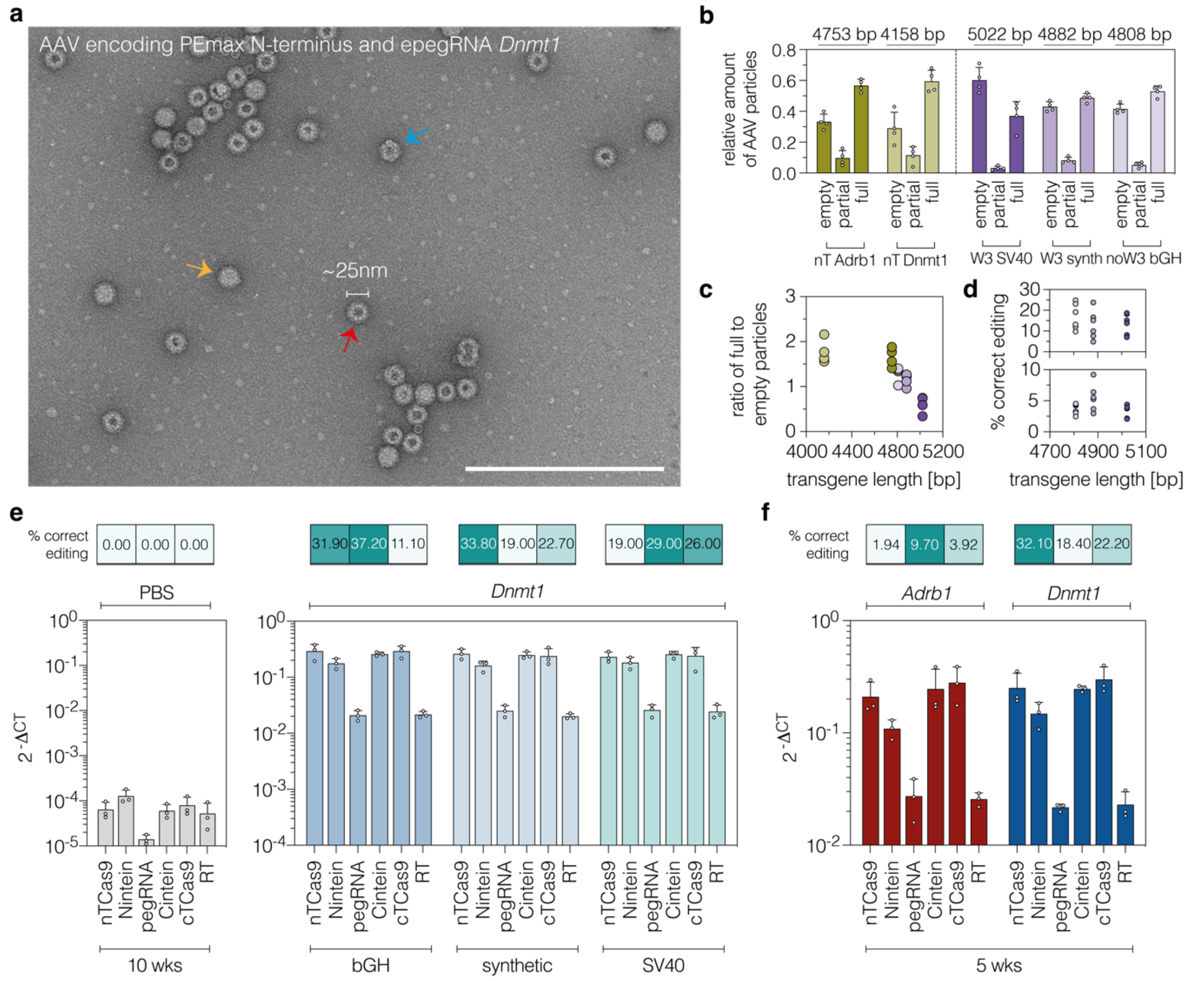


**Extended data figure 6: Detailed *in vitro* and *in vivo* analyses of AAV preparations. (a)** Representative electron micrograph of a negative-stained AAV preparation (AAV-PEmax-nT with the epegRNA targeting *Dnmt1*) containing fully-packaged (orange arrow), partially-packaged (blue arrow), and empty particles (red arrow). Scale bar, 250 nm. (**b-d**) Relationship between transgene length and packaging efficiency (b,c) or editing efficiency (d) at the *Dnmt1* and *Adrb1* locus. Transgene lengths in base pairs (bp) are indicated at the top above the respective construct (b). The color coding of AAV preparations in (c) and (d) corresponds to the same colors as in (b). (**e,f**) PEmax and epegRNA transcript levels of AAV-nT and -cT preparations targeting *Dnmt1* (e,f) or *Adrb1* (f) in mouse striata. Corresponding editing rates were determined by deep amplicon sequencing and are depicted for every animal above each construct (e,f). Transcript levels were normalized to *Gapdh*. Data are represented as means±s.d. of four individual grid squares (a-c) or three animals per group (d-f). nT/cT, N-/C-terminal PEmax AAV vector; wks, weeks.

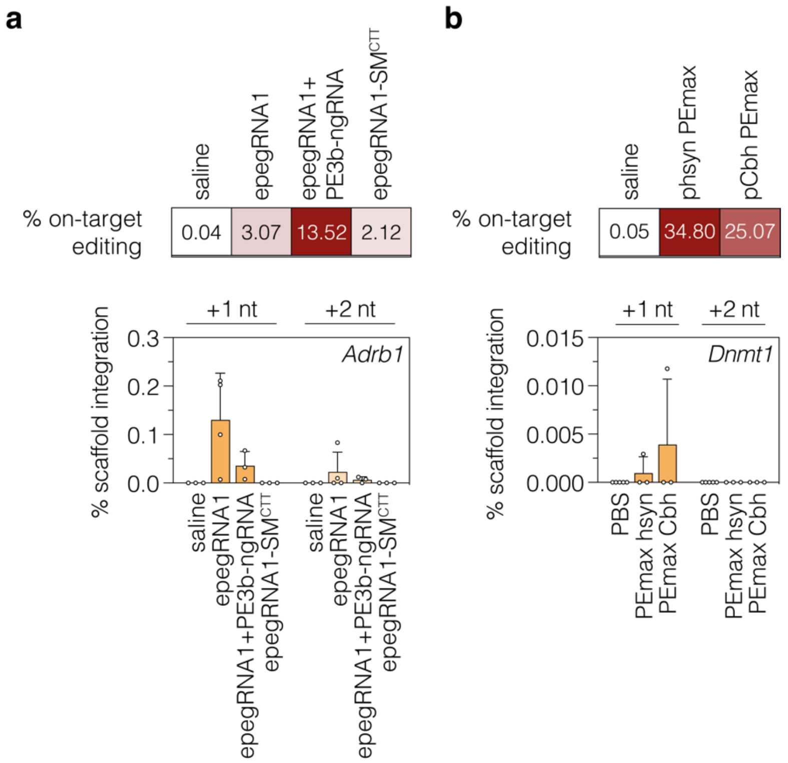


**Extended data figure 7: Scaffold integrations are low at the *Adrb1* and *Dnmt1* locus.** (a,b) Integration of the first (+1) and second (+2) nt of the sgRNA scaffold into the genome at the *Adrb1* and *Dnmt1* locus. Average on-target editing rates of the respective animals are displayed on top. Data are represented as means±s.d. of at least three animals. Each data point represents one mouse.

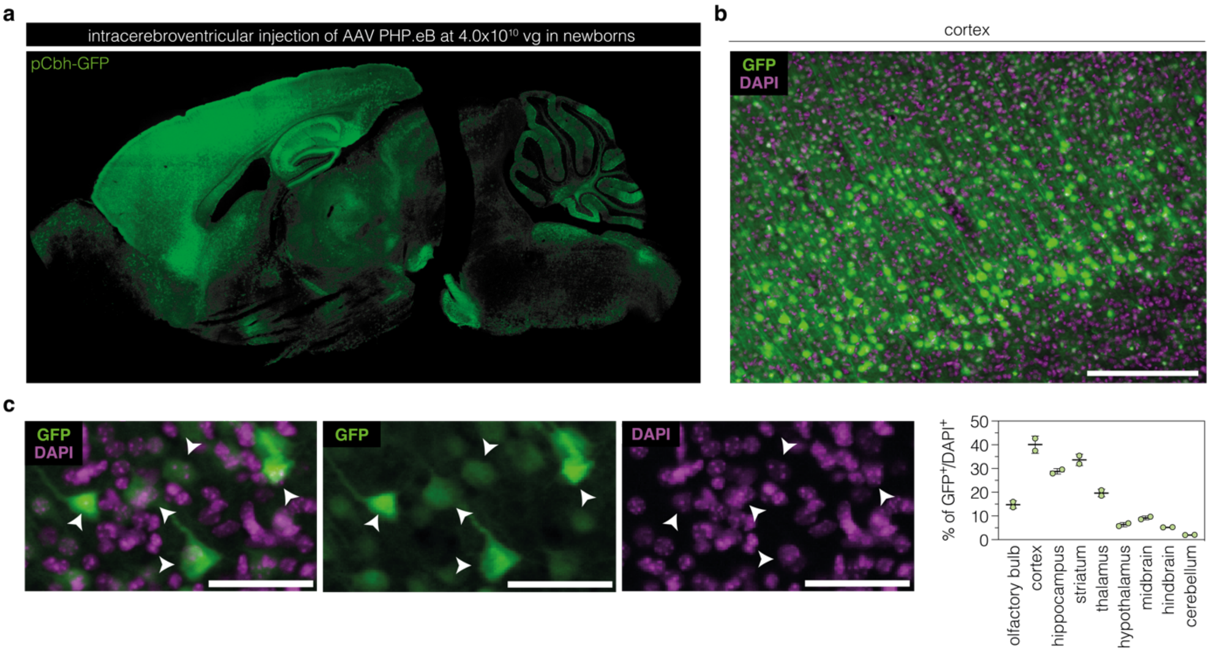


**Extended data figure 8: Transduction efficiency of AAV serotype PHP.eB after ICV injection into neonatal mice.** (**a**) Representative whole-brain image of GFP fluorescence after 30 days of expression. AAV particles, expressing EGFP under the Cbh promoter, were delivered to newborn mice at a dose of $4\times10^{10}$ vg per animal via ICV injection. (**b,c**) Representative image of GFP transgene expression (green) and DAPI staining (magenta) in the cortex and quantification of DAPI$^+$ cells that were transduced by the AAV in different brain regions. Data are displayed as mean±range (n=2 mice). Scale bars, 200 µm (b) and 50 µm (c). vg, vector genomes; GFP, green fluorescent protein.

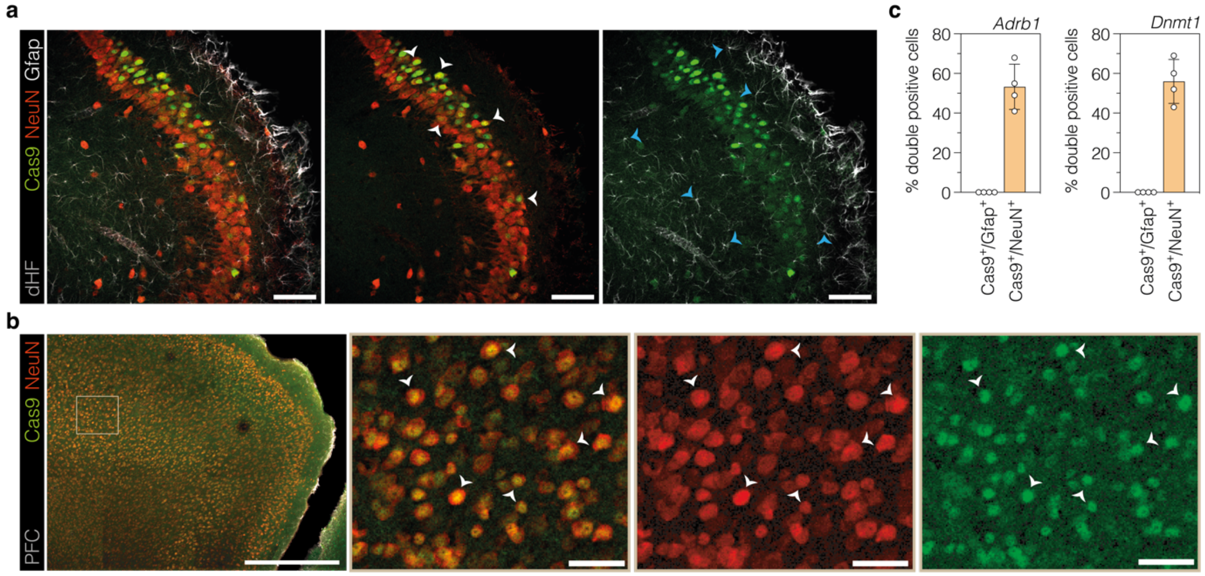


**Extended data figure 9: Neuron-specific Cas9 expression in the mouse hippocampus and prefrontal cortex.** (**a,b**) Representative fluorescence micrographs of *Sp*Cas9, NeuN, and Gfap in the hippocampus (a) and *Sp*Cas9 and NeuN in the prefrontal cortex (b) of treated newborn mice at 10 weeks post-injection (hsyn-PE3bmax targeting the *Adrb1* locus). (**c**) Quantifications of the overlap between *Sp*Cas9 expression (Cas9, green) in neurons (NeuN, white arrowheads) and astrocytes (Gfap, blue arrowheads) in mice treated with hsyn-PEmax targeting *Dnmt1* or hsyn-PE3bmax targeting *Adrb1*. Data are represented as means±s.d. of four independently imaged tissue regions (one mouse per locus).

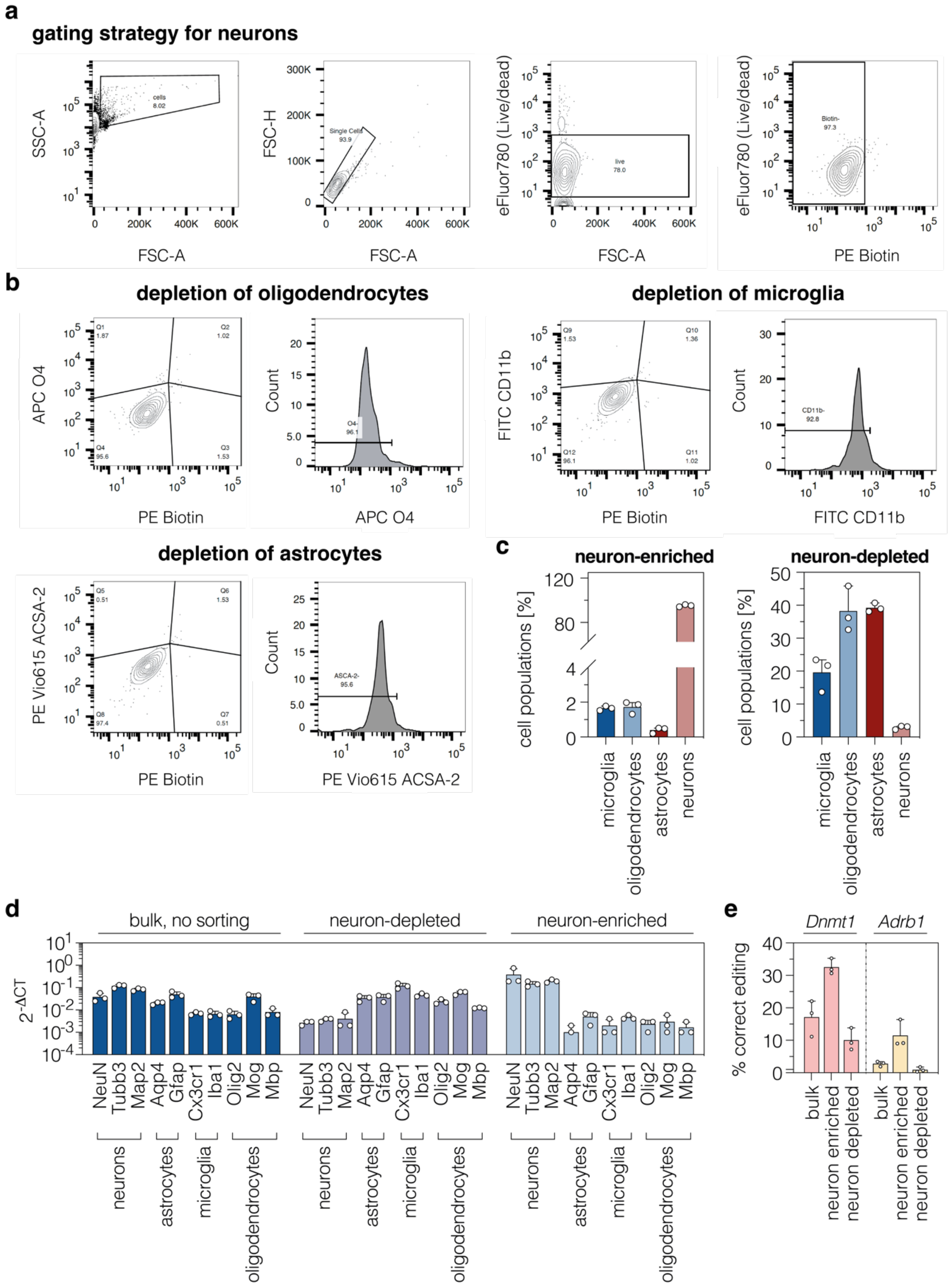


**Extended data figure 10: Validation of MACS-purified neuronal populations.** (**a-c**) Gating strategy for validation of oligodendrocyte-, microglia-, and astrocyte-depletion in neuron-enriched samples (a,b) and quantification of the respective populations (c). (**d**) Transcript levels of neuronal, astrocytic, microglial, and oligodendrocytic markers in bulk, neuron-depleted, and neuron-enriched samples. Transcript levels were normalized to *Gapdh*. Bulk samples are defined as single-cell suspensions before magnetic sorting. (**e**) Relative neuron-enrichment and

-depletion at the *Adrb1* and *Dnmt1* locus (hsyn-PEmax-*Dnmt1* or hsyn-PE3bmax-*Adrb1* at 10 weeks post-injection). Data are represented as means±s.d. of three animals (per locus).

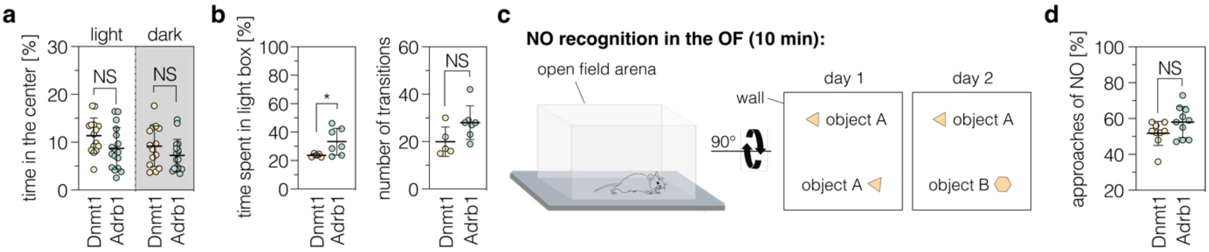


**Extended data figure 11: Anxiety-related behaviors and memory performance in newborn mice.** (**a,b**) Time spent in the center of the OF arena (a) or in the light compartment in the LD transition test (b) for Dnmt1- and Adrb1-treated newborns (n=5-16 mice per group). (**c**) Schematic representation of the novel object (NO) recognition test. The duration of the test is indicated in brackets. (**d**) Percentage of approaches of the NO for Dnmt1- and Adrb1-treated newborns (n=9-10 mice per group). Adrb1- and Dnmt1-injected mice were kept in a 12:12 light/dark cycle; areas highlighted in gray indicate the dark cycle. Data are displayed as means±s.d. and were analyzed using a two-tailed Student's *t*-test with Welch's correction (NS, $P>0.05$; $**P<0.005$). Each data point represents one animal. NS, not significant.

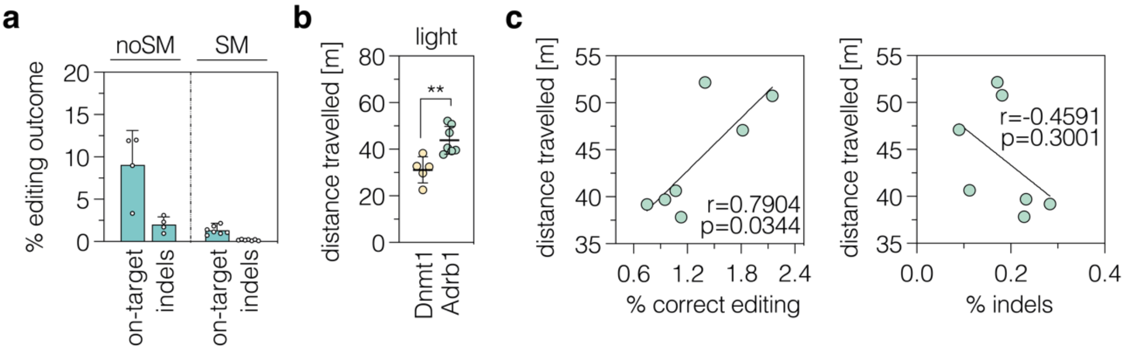


**Extended data figure 12: Behavioral changes are related to on-target editing and not indels.** (**a**) Endpoint on-target editing and indel rates in cortices of newborn mice injected with either epegRNA1/PE3bmax (no SM, n=4 mice) or epegRNA1-SM$^{CTT}$/PEmax (SM, n=7 mice). (**b,c)** Locomotor activity in the OF (b) and Pearson correlations of correct editing or indels and distance traveled (c) for epegRNA1-SM$^{CTT}$-treated animals (n=5-7 mice per group). Correlation coefficients and p-values are indicated in the respective plots. Adrb1- and Dnmt1-injected mice were kept in a 12:12 light/dark cycle; areas highlighted in gray indicate the dark cycle. Data are displayed as means±s.d. and were analyzed using a two-tailed Student's *t*-test with Welch's correction (**P<0.05*; ***P<0.005*). Each data point represents one animal.

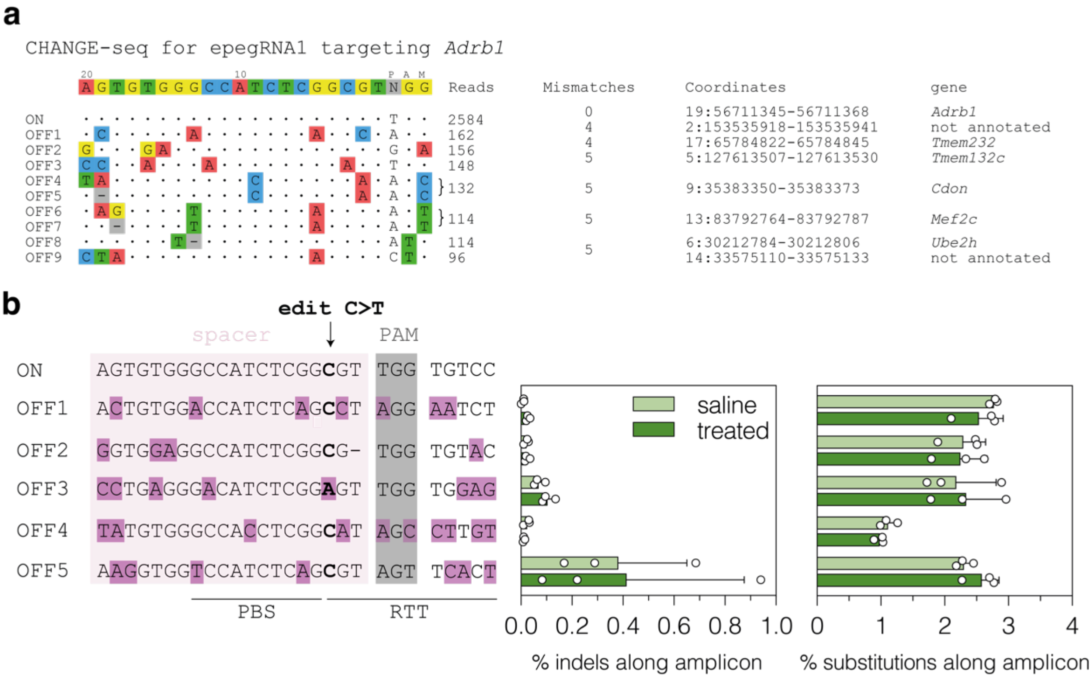


**Extended data figure 13: *In vivo* prime editing does not induce off-target editing.** (**a**) Off-targets for the pegRNA protospacer targeting the Adrb1 locus were experimentally identified by CHANGE-seq[43]. Off-targets with <90 reads are not shown. (**b**) Deep sequencing of the top 5 off-targets, identified by CHANGE-seq, in saline- and PE-treated mice (>15'000 reads per site, n=3 mice per group). The spacer sequence (rosé), mismatches (purple), PAM sequence (gray), and on-target edit (bold) are indicated. Indels and bystanders were quantified across the whole amplicon length. Data are displayed as means ±s.d. and were analyzed using a two-tailed Student's t-test- If not indicated otherwise, differences were not significant.

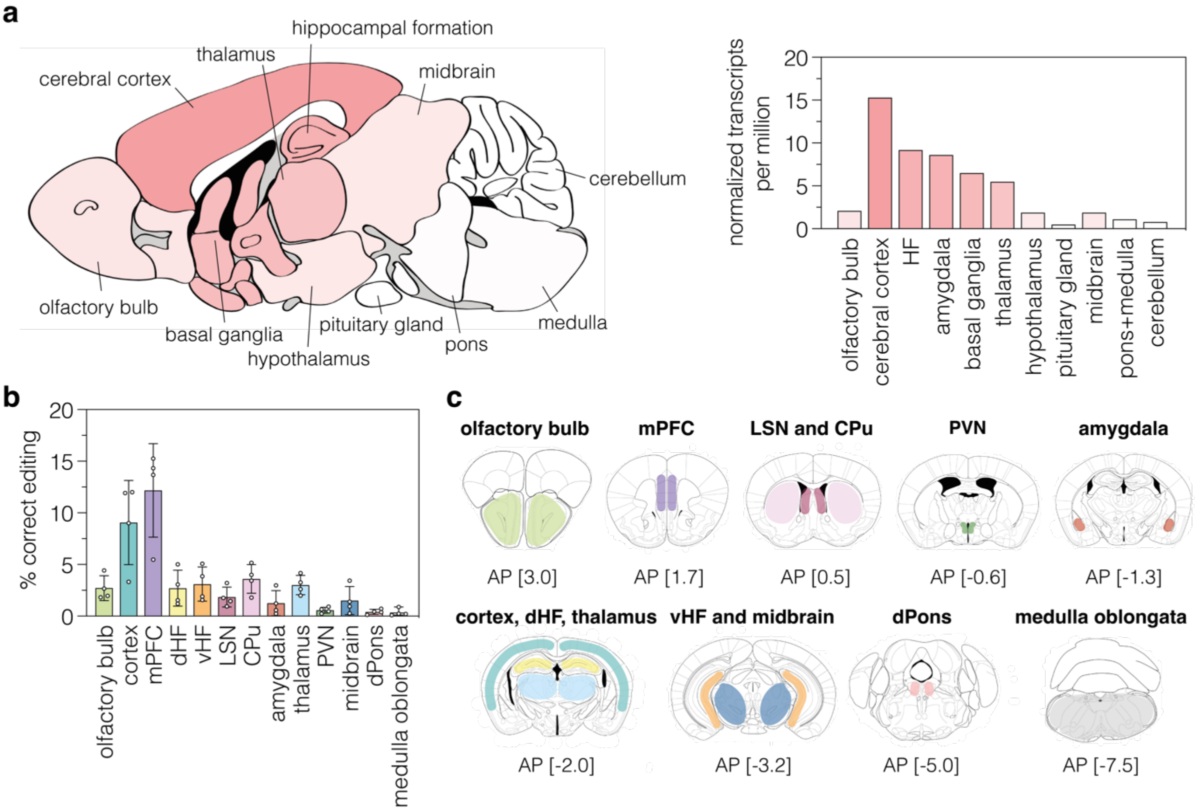


**Extended data figure 14: Prime editing across *Adrb1*-expressing brain regions.** (**a**,**b**) Color-coded *Adrb1*-expression (a) and normalized transcripts per million (nTPM, b) across the mouse brain (image credit: Human protein atlas[46]). Darker shades indicate higher expression. (**c**) Editing rates in brain regions with *Adrb1* expression (cut-off >1.0 nTPM in the mouse protein atlas[46]) in mice injected as newborns (n=4 mice; hsyn-PE3bmax). Each data point represents one animal. (**d**) The location of each region that was isolated for deep sequencing is indicated on the mouse brain atlas[49]. mPFC, medial prefrontal cortex; LSN, lateral septal nucleus; CPu, caudate putamen; PVN, paraventricular nucleus; d/vHF, dorsal/ventral hippocampal formation; dPons, dorsal pons. Color coding of isolated brain areas is identical in (c) and (d).

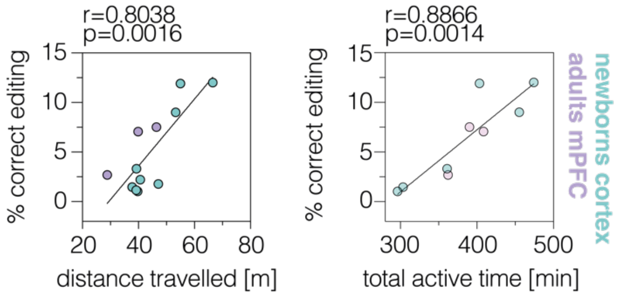


**Extended data figure 15: Correlation of prime editing rates and locomotion or general activity in newborn and adult mice.** Pearson correlation of prime editing frequency and distance traveled in the OF (left, dark cycle) or total active time in the home cage (right, dark cycle). Newborn mice are labelled in green; adult mice are highlighted in purple. Each data point represents one animal.

### Extended data tables

Extended data table 1: Oligos used for cloning of PE and AAV plasmids.

| oligo name | oligo sequence (5’→ 3’) |
|---|---|
| N-intein_fwd | CAGATCCGCTAGAGATCCGCGGCCGCTAATACGACTCACTATAGGGAGAGCC |
| C-intein_rev | TCCTCTTCTTCTTGGGCTCGAATTCGCTGCCGTCGGCGGTTCT |
| split867_fwd | GCAGCGGTGGCGGCGGCAGCAGCGACAACGTGCCCTCC |
| split866_rev | CCGCCACCGCTGCCGCCGCCACCCTTGCCCCGGTTCTTGTCG |
| split1025_fwd | GCAGCGGTGGCGGCGGCAGCAGCGAGCAGGAAATCGGCAAG |
| split1024_rev | CCACCGCTGCCGCCGCCACCCTTGGCGATCATCTTCCG |
| split573_fwd | GCAGCGGTGGCGGCGGCAGCTGCTTCGACTCCGTGGAAATCTCC |
| split572_rev | CCGCCACCGCTGCCGCCGCCACCCTCGATTTCTTGAAGTAGTCCTC |
| split713_fwd | GCAGCGGTGGCGGCGGCAGCTCCGGCCAGGGCGATA |
| split712_rev | CCGCCACCGCTGCCGCCGCCACCCACCTGGGCTTTCTGGATGT |
| AAV-BPNLS-PEmax712_fwd | GAGCGCAGTCGAGAGTTGGACCGGTGCCACCATGAAACGGACAG |
| AAV-BPNLS-PEmax713_rev | TCTGTTGGCGAAGCCGTCGGACTTCAGGAAATCCAGGATTG |
| AAV-BPNLS-Nintein_fwd | AAGCGGAAAGTCGACAAGAAGTACAGCATC |
| AAV-BPNLS-Nintein_rev | CGCCAGAATTAGGCAGGTTATCCACTCTCA |
| AAV-pegRNA-optscf_fwd | ACTCCATCACTAGGGGTTCCTGCGGCCGCGAGGGCCTATTTCCCATG |
| AAV-pegRNA-optscf_rev | GATGCGGTGGGCTCTATGGCTCGAGAAAAAAAGCCATCTCGGTG |
| AAV-pegRNAevo-optscf_rev | GATGCGGTGGGCTCTATGGCTCGAGAAAAAAATTCTAGTTGGTTTAACGC |
| AAV-3bpegRNAevo-optscf_rev | ATCATGGGAAATAGGCCCTCGAATTCAAAAAAATTCTAGTTGGTTTAACGC |
| AAV-3bpegRNAevo-optscf_fwd | AAACCAACTAGAATTTTTTTGAATTCGAGGGCCTATTTCCCATG |
| AAV-3bngRNA-optscf_rev | GATGCGGTGGGCTCTATGGCTCGAGAAAAAAAAGCACCGACTC |
| AAV-3bngRNAevo-optscf_rev | GATGCGGTGGGCTCTATGGCTCGAGAAAAAAAAATTCTAGTTGGTTTAACGC |
| pegRNA-U6-NdeI_fwd | TGTTTTAAAATGGACTATCATATGCTTACCGTAACTTGAAAGTATTTC |
| AAV-bGHpolyA_fwd | GATGCGGTGGGCTCTATGG |
| AAV-U6_rev | GGCCGCGAGGGCCTATTT |

Extended data table 2: Oligos used for cloning of pegRNA and nicking sgRNA plasmids.

| oligo name | oligo sequence (5’→ 3’) |
|---|---|
| pegRNA1-Adrb1-spacer_fwd | CACCGAGTGTGGGCCATCTCGGCGTGTTTT |
| pegRNA1-Adrb1-spacer_rev | CTCTAAAACACGCCGAGATGGCCCACACTC |
| pegRNA1.1-Adrb1-ext_fwd | GTGCGGACACCAACACCGAGATGGC |
| pegRNA1.1-Adrb1-ext_rev | AAAAGCCATCTCGGTGTTGGTGTCC |
| pegRNA1.2-Adrb1-ext_fwd | GTGCAAGGACACCAACACCGAGATGGC |
| pegRNA1.2-Adrb1-ext_rev | AAAAGCCATCTCGGTGTTGGTGTCCTT |
| pegRNA1.3-Adrb1-ext_fwd | GTGCGGAAGGACACCAACACCGAGATGGC |
| pegRNA1.3-Adrb1-ext_rev | AAAAGCCATCTCGGTGTTGGTGTCCTTCC |
| pegRNA1.4-Adrb1-ext_fwd | GTGCGGACACCAACACCGAGATGGCCCA |
| pegRNA1.4-Adrb1-ext_rev | AAAATGGGCCATCTCGGTGTTGGTGTCC |
| pegRNA1.5-Adrb1-ext_fwd | GTGCAAGGACACCAACACCGAGATGGCCCA |
| pegRNA1.5-Adrb1-ext_rev | AAAATGGGCCATCTCGGTGTTGGTGTCCTT |
| pegRNA1.6-Adrb1-ext_fwd | GTGCGGAAGGACACCAACACCGAGATGGCCCA |
| pegRNA1.6-Adrb1-ext_rev | AAAATGGGCCATCTCGGTGTTGGTGTCCTTCC |
| pegRNA-scaffold_fwd | AGAGCTAGAAATAGCAAGTTAAAATAAGGCTAGTCCGTTATCAACTTGAAAAAGTGGCACCGAGTCG |
| pegRNA-scaffold_rev | GCACCGACTCGGTGCCACTTTTTCAAGTTGATAACGGACTAGCCTTATTTTAACTTGCTATTTCTAG |
| GG-epegRNA1-spacer_fwd | CACCGAGTGTGGGCCATCTCGGCGTGTTTC |
| GG-epegRNA1-spacer_rev | CTCTGAAACACGCCGAGATGGCCCACACTC |
| GG-scaffold-optscf_fwd | AGAGCTATGCTGGAAACAGCATAGCAAGTTGAAATAAGGCTAGTCCGTTATCAACTTGAAAAAGTGGCACCGAGTCG |
| GG-scaffold-optscf_rev | GCACCGACTCGGTGCCACTTTTTCAAGTTGATAACGGACTAGCCTTATTTCAACTTGCTATGCTGTTTCCAGCATAG |
| GG-evoRT/PBS-SM1_fwd | GTGCGGACACCAGCACCGAGATGGC |
| GG-evoRT/PBS-SM1_rev | CGCGGCCATCTCGGTGCTGGTGTCC |
| GG-evoRT/PBS-SM2_fwd | GTGCGGACACTAACACCGAGATGGC |
| GG-evoRT/PBS-SM2_rev | CGCGGCCATCTCGGTGTTAGTGTCC |
| GG-evoRT/PBS-SM3_fwd | GTGCGGACACAAGCACCGAGATGGC |
| GG-evoRT/PBS-SM3_rev | CGCGGCCATCTCGGTGCTTGTGTCC |
| GG-evoRT/PBS-SM4_fwd | GTGCGGACACGAGCACCGAGATGGC |

| Extended data table 2 continued. | |
|---|---|
| oligo name | oligo sequence (5’→ 3’) |
| GG-evoRT/PBS-SM4_rev | CGCGGCCATCTCGGTGCTCGTGTCC |
| GG-evoRT/PBS-SM5_fwd | GTGCGGACACTAGCACCGAGATGGC |
| GG-evoRT/PBS-SM5_rev | CGCGGCCATCTCGGTGCTAGTGTCC |
| PE3ngRNA3.1_fwd | CCACGGCCCACACTGTGCACACGA |
| PE3ngRNA3.1_rev | AAACTCGTGTGCACAGTGTGGGCC |
| PE3ngRNA3.2_fwd | CACCGAAGGACACCAACGCCGAGA |
| PE3ngRNA3.2_rev | AAACTCTCGGCGTTGGTGTCCTTC |
| PE3ngRNA3.3_fwd | CCACGCGACGTGATGGCGAGGTAG |
| PE3ngRNA3.3_rev | AAACCTACCTCGCCATCACGTCGC |
| PE3ngRNA3.4_fwd | CCACGCGCGCGTCAGCAAACTC |
| PE3ngRNA3.4_rev | AAACGAGTTTGCTGACGCGCGCGC |
| U6-epegRNAs_fwd | TGTGGAAAGGACGAAACACC |
| GG-epeg-tmpknot_rev | CAGGGGGGGGCTCCTGACCCTGTACTGTGTGCAGTGGaAG |
| GG-epeg-evopreQ1_rev | TAACGCGTAACTAGATAGAACCGCGTGTACTGTGTGCAGTGGaAG |
| Dnmt1_spacer-fwd | caccGCGGGCTGGAGCTGTTCGCGCgtttt |
| Dnmt1_spacer-rev | ctctaaaacGCCTAATGTACTGTGTGCAGc |
| Dnmt1_RT/PBS-fwd | gtgcAAGATGgCAGCGCGAACAGCTCCAG |
| Dnmt1_RT/PBS-rev | aaaaCTGGAGCTGTTCGCGCTGCCATCTT |
| Adrb1-epeg-tmpknot_rev | CAGGGGGGGGCTCCTGACCCGCCATCTCGGTGTTGGTGTC |
| Adrb1-epeg-evopreQ1_rev | TAACGCGTAACTAGATAGAACCGCGGCCATCTCGGTGTTGGTGTC |
| PE3bngRNA-Adrb1-tmp_rev | CAGGGGGGGGCTCCTGACCCGCACCGACTCGGTGCCACTT |
| PE3bngRNA-Adrb1-evo_rev | TAACGCGTAACTAGATAGAACCGCGGCACCGACTCGGTGCCACTT |
| pegRNA-eIF2B4_spacer_fwd | CACCGAGACCAGATTCAACA ACCAGGTTTT |
| pegRNA-eIF2B4_spacer_rev | CTCTAAAACCTGGTTGTTGAATCTGGTCTC |
| pegRNA-eIF2B4_RT/PBS_fwd | GTGCGTCCCTCCGGTTGTTGAATCTG |
| pegRNA-eIF2B4_RT/PBS_rev | AAAACAGATTCAACAACCGGAGGGAC |
| evo-eIF2B4_rev | CCCAAGCTTAAAAAAATTCTAGTTGGTTTAACGCGTAACTAGATAGAA CCGCGCAGATTCAACAACCGGA |
| PE3ngRNA-eIF2B4_fwd | CACCGCAGTGTAAGCGGGGAGACC |
| PE3ngRNA-eIF2B4_rev | AAACGGTCTCCCCGCTTACACTGC |
| evo-ngRNAs_rev | ACGCCAAGCTTAAAAAAATTCTAGTTGGTTTAACGCGTAACTAGATAGA ACCGCGGCACCGACTCGGTGC |
| pegRNA-eIF2B5_spacer_fwd | CACCGCCATCACCACGTTGTCCTCAGTTTT |
| pegRNA-eIF2B5_spacer_rev | CTCTAAAACTGAGGACAACGTGGTGATGGC |
| pegRNA-eIF2B5_RT/PBS_fwd | GTGCACACGCTGCCATGAGGACAACGT |
| pegRNA-eIF2B5_RT/PBS_rev | AAAAACGTTGTCCTCATGGCAGCGTGT |
| evo-eIF2B5_rev | CCCAAGCTTAAAAAAATTCTAGTTGGTTTAACGCGTAACTAGATAGAA CCGCGACGTTGTCCTCATGGC |
| PE3ngRNA-eIF2B5_fwd | CACCGAAATGTCTCTGTGATGACAA |
| PE3ngRNA-eIF2B5_rev | AAACTTGTCATCACAGAGACATTTC |
| pegRNA-otc_spacer_fwd | CACCGACCACACAAGACATTCACTTGTTTT |
| pegRNA-otc_spacer_rev | CTCTAAAACAAGTGAATGTCTTGTGTGGTC |
| pegRNA-otc_RT/PBS_fwd | GTGCACCGAGCGGTGTCTGTGAGACTTTCATTCACACCCAAGTGAATG TCTTGTG |
| pegRNA-otc_RT/PBS_rev | AAAACACAAGACATTCACTTGGGTGTGAATGAAAGTCTCACAGACACC GCTCGGT |
| evo-otc_rev | CCCAAGCTTAAAAAAATTCTAGTTGGTTTAACGCGTAACTAGATAGAA CCGCGCACAAGACATTCACTTGGGT |
| PE3ngRNA-otc_fwd | CACCGACCACACAAGACATTCACTT |
| PE3ngRNA-otc_rev | AAACAAGTGAATGTCTTGTGTGGTC |

Extended data table 3: Oligos used for cloning of shRNAs.

| oligo name | oligo sequence (5’→ 3’) |
|---|---|
| MLH1humanshRNA-1_fwd | CCGGAAGTTGATTCAGATCCAAGACTCGAGTCTTGGATCTGAATCAACTTTT TTTG |
| MLH1humanshRNA-3_fwd | CCGGCCAAGTGAAGAATATGGGAAACTCGAGTTTCCCATATTCTTCACTTGG TTTTTG |
| MLH1humanshRNA-5_fwd | CCGGAATCCACAAGTATTCAAGTGACTCGAGTCACTTGAATACTTGTGGATT TTTTTG |
| MLH1humanshRNA-1_rev | AATTCAAAAAAAGTTGATTCAGATCCAAGACTCGAGTCTTGGATCTGAATCA ACTT |
| MLH1humanshRNA-3_rev | AATTCAAAAACCAAGTGAAGAATATGGGAAACTCGAGTTTCCCATATTCTTC ACTTGG |
| MLH1humanshRNA-5_rev | AATTCAAAAAAATCCACAAGTATTCAAGTGACTCGAGTCACTTGAATACTTG TGGATT |
| MLH1mouseshRNA-2_fwd | CCGGCCGAAGCATTTCACAGAAGATCTCGAGATCTTCTGTGAAATGCTTCGG TTTTTG |

Extended data table 3 continued.

| oligo name | oligo sequence (5'→ 3') |
| --- | --- |
| MLH1mouseshRNA-3_fwd | CCGGGCTAATTCAGATCCAAGACAACTCGAGTTGTCTTGGATCTGAATTAGCTTTTTG |
| MLH1mouseshRNA-4_fwd | CCGGAATCTACAAATATTCAAGTGGCTCGAGCCACTTGAATATTTGTAGATTTTTTTG |
| MLH1mouseshRNA-2_rev | AATTCAAAAACCGAAGCATTTCACAGAAGATCTCGAGATCTTCTGTGAAATGCTTCGG |
| MLH1mouseshRNA-3_rev | AATTCAAAAAGCTAATTCAGATCCAAGACAACTCGAGTTGTCTTGGATCTGAATTAGC |
| MLH1mouseshRNA-4_rev | AATTCAAAAAAATCTACAAATATTCAAGTGGCTCGAGCCACTTGAATATTTGTAGATT |
| MSH2humanshRNA-1_fwd | CCGGATTCATGTTGCAGAGCTTGCTCTCGAGAGCAAGCTCTGCAACATGAATTTTTTG |
| MSH2humanshRNA-3_fwd | CCGGGCCTTGCTGAATAAGTGTAAACTCGAGTTTACACTTATTCAGCAAGGCTTTTTG |
| MSH2humanshRNA-4_fwd | CCGGAAGACATTCTCTTTGGTAACACTCGAGTGTTACCAAAGAGAATGTCTTTTTTTG |
| MSH2humanshRNA-1_rev | AATTCAAAAAATTCATGTTGCAGAGCTTGCTCTCGAGAGCAAGCTCTGCAACATGAAT |
| MSH2humanshRNA-3_rev | AATTCAAAAAGCCTTGCTGAATAAGTGTAAACTCGAGTTTACACTTATTCAGCAAGGC |
| MSH2humanshRNA-4_rev | AATTCAAAAAAAGACATTCTCTTTGGTAACACTCGAGTGTTACCAAAGAGAATGTCTT |
| MSH2mouseshRNA-1_fwd | CCGGCGAGATCATTTCACGGATAAACTCGAGTTTATCCGTGAAATGATCTCGTTTTTG |
| MSH2mouseshRNA-3_fwd | CCGGAACTTTGAGTCTTTCGTGAAACTCGAGTTTCACGAAAGACTCAAAGTTTTTTTG |
| MSH2mouseshRNA-4_fwd | CCGGAAGCTGGAAATAAGGCGTCTACTCGAGTAGACGCCTTATTTCCAGCTTTTTTTG |
| MSH2mouseshRNA-1_rev | AATTCAAAAACGAGATCATTTCACGGATAAACTCGAGTTTATCCGTGAAATGATCTCG |
| MSH2mouseshRNA-3_rev | AATTCAAAAAAACTTTGAGTCTTTCGTGAAACTCGAGTTTCACGAAAGACTCAAAGTT |
| MSH2mouseshRNA-4_rev | AATTCAAAAAAAGCTGGAAATAAGGCGTCTACTCGAGTAGACGCCTTATTTCCAGCTT |

Extended data table 4: Oligos used for RT-qPCR.

| oligo name | oligo sequence (5'→ 3') |
| --- | --- |
| RTqPCR-mAdrb1_fwd | GCTCTGGACTTCGGTAGATGTG |
| RTqPCR-mAdrb1_rev | CGTCAGCAAACTCTGGTAGCGA |
| RTqPCR-hAdrb1_fwd | TTCCTGCCCATCCTCATGCACT |
| RTqPCR-hAdrb1_rev | GTAGAAGGAGACTACGGACGAG |
| RTqPCR-hMSH2_fwd | CAGCAGTCAGAGCCCTTAACCT |
| RTqPCR-hMSH2_rev | GAGAGGCTGCTTAATCCACTGG |
| RTqPCR-mMSH2_fwd | GAACAAAGGCGAGTATGAAGAGG |
| RTqPCR-mMSH2_rev | GCGTCTAAGTGAGCCAGCACAT |
| RTqPCR-hMLH1_fwd | GCCTTGGCACAGCATCAAACCA |
| RTqPCR-hMLH1_rev | ATGGCAAGGTCAAAGAGCGGT |
| RTqPCR-mMLH1_fwd | CCTCCAGGATGTATTTCACCCAG |
| RTqPCR-mMLH1_rev | ACGGACCATCTGGTAAGCGTAG |
| RTqPCR-hactB_fwd | CACCATTGGCAATGAGCGGTTC |
| RTqPCR-hactB_rev | AGGTCTTTGCGGATGTCCACGT |
| RTqPCR-mactB_fwd | GGCTGTATTCCCCTCCATCG |
| RTqPCR-mactB_rev | CCAGTTGGTAACAATGCCATGT |
| RTqPCR-hGAPDH_fwd | GTCTCCTCTGACTTCAACAGCG |
| RTqPCR-hGAPDH_rev | ACCACCCTGTTGCTGTAGCCAA |
| RTqPCR-mGAPDH_fwd | CATCACTGCCACCCAGAAGACTG |
| RTqPCR-mGAPDH_rev | ATGCCAGTGAGCTTCCCGTTCAG |
| RTqPCR-NeuN_fwd | CACCACTCTCTTGTCCGTTTGC |
| RTqPCR-NeuN_rev | GGCTGAGCATATCTGTAAGCTGC |
| RTqPCR-Tubb3_fwd | CATCAGCGATGAGCACGGCATA |
| RTqPCR-Tubb3_rev | GGTTCCAAGTCCACCAGAATGG |
| RTqPCR-Map2_fwd | GCTGTAGCAGTCCTGAAAGGTG |
| RTqPCR-Map2_rev | CTTCCTCCACTGTGGCTGTTTG |
| RTqPCR-Aqp4_fwd | AGCCAGCATGAATCCAGCTCGA |
| RTqPCR-Aqp4_rev | TCATAAAGGGCACCTGCCAGCA |
| RTqPCR-Gfap_fwd | ACATCGAGATCGCCACCTACA |
| RTqPCR-Gfap_rev | CCACGATGTTCCTCTTGAGGTG |

| Extended data table 4 continued. | |
|---|---|
| oligo name | oligo sequence (5'→ 3') |
| RTqPCR-Iba1_fwd | TCTGCCGTCCAAACTTGAAGCC |
| RTqPCR-Iba1_rev | CTCTTCAGCTCTAGGTGGGTCT |
| RTqPCR-Cx3cr1_fwd | GAGCATCACTGACATCTACCTCC |
| RTqPCR-Cx3cr1_rev | AGAAGGCAGTCGTGAGCTTGCA |
| RTqPCR-Olig2_fwd | ATGCACGACCTCAACATCGCCA |
| RTqPCR-Olig2_rev | ACCAGTCGCTTCATCTCCTCCA |
| RTqPCR-Mog_fwd | GATGAAGGAGGCTACACCTGCT |
| RTqPCR-Mog_rev | CGTAGGCACAAGTGCGATGAGA |
| RTqPCR-Mbp_fwd | ATTCACCGAGGAGAGGCTGGAA |
| RTqPCR-Mbp_rev | TGTGTGCTTGGAGTCTGTCACC |

Extended data table 5: Amino acid sequences of intein-split PEmax p.713 and p.714 constructs.

Intein-split PEmax p.713: p.NLS/n*Sp*Cas9$^{1-713}$(R221K,N394K)/linker/N-intein/NLS

MKRTADGSEFESPKKKRKVDKKYSIGLDIGTNSVGWAVITDEYKVPSKKFKVLGNTDRHSIKKNLIG
ALLFDSGETAEATRLKRTARRRYTRRKNRICYLQEIFSNEMAKVDDSFFHRLEESFLVEEDKKHERH
PIFGNIVDEVAYHEKYPTIYHLRKKLVDSTDKADLRLIYLALAHMIKFRGHFLIEGDLNPDNSDVDKL
FIQLVQTYNQLFEENPINASGVDAKAILSARLSKSRKLENLIAQLPGEKKNGLFGNLIALSLGLTPNFK
SNFDLAEDAKLQLSKDTYDDDLDNLLAQIGDQYADLFLAAKNLSDAILLSDILRVNTEITKAPLSAS
MIKRYDEHHQDLTLLKALVRQQLPEKYKEIFFDQSKNGYAGYIDGGASQEEFYKFIKPILEKMDGTE
ELLVKLKREDLLRKQRTFDNGSIPHQIHLGELHAILRRQEDFYPFLKDNREKIEKILTFRIPYYVGPLA
RGNSRFAWMTRKSEETITPWNFEEVVDKGASAQSFIERMTNFDKNLPNEKVLPKHSLLYEYFTVYN
ELTKVKYVTEGMRKPAFLSGEQKKAIVDLLFKTNRKVTVKQLKEDYFKKIECFDSVEISGVEDRFNA
SLGTYHDLLKIIKDKDFLDNEENEDILEDIVLTLTLFEDREMIEERLKTYAHLFDDKVMKQLKRRRYT
GWGRLSRKLINGIRDKQSGKTILDFLKSDGFANRNFMQLIHDDSLTFKEDIQKAQVGGGGSGGGGSG
GGGSCLSYETEILTVEYGLLPIGKIVEKRIECTVYSVDNNGNIYTQPVAQWHDRGEQEVFEYCLEDGS
LIRATKDHKFMTVDGQMLPIDEIFERELDLMRVDNLPNSGGSKRTADGSEFEPKKKRKV*

Intein-split PEmax p.714: p.NLS/C-intein/linker/n*Sp*Cas9$^{714-1368}$(H840A)/linker-NLS-linker/RT-dRnH/NLS

MKRTADGSEFESPKKKRKVIKIATRKYLGKQNVYDIGVERDHNFALKNGFIASNGGGGSGGGGSGG
GGSSGQGDSLHEHIANLAGSPAIKKGILQTVKVVDELVKVMGRHKPENIVIEMARENQTTQKGQKN
SRERMKRIEEGIKELGSQILKEHPVENTQLQNEKLYLYYLQNGRDMYVDQELDINRLSDYDVDAIVP
QSFLKDDSIDNKVLTRSDKNRGKSDNVPSEEVVKKMKNYWRQLLNAKLITQRKFDNLTKAERGGL
SELDKAGFIKRQLVETRQITKHVAQILDSRMNTKYDENDKLIREVKVITLKSKLVSDFRKDFQFYKV
REINNYHHAHDAYLNAVVGTALIKKYPKLESEFVYGDYKVYDVRKMIAKSEQEIGKATAKYFFYSN
IMNFFKTEITLANGEIRKRPLIETNGETGEIVWDKGRDFATVRKVLSMPQVNIVKKTEVQTGGFSKES
ILPKRNSDKLIARKKDWDPKKYGGFDSPTVAYSVLVVAKVEKGKSKKLKSVKELLGITIMERSSFEK
NPIDFLEAKGYKEVKKDLIIKLPKYSLFELENGRKRMLASAGELQKGNELALPSKYVNFLYLASHYE
KLKGSPEDNEQKQLFVEQHKHYLDEIIEQISEFSKRVILADANLDKVLSAYNKHRDKPIREQAENIIHL
FTLTNLGAPAAFKYFDTTIDRKRYTSTKEVLDATLIHQSITGLYETRIDLSQLGGDSGGSSGGSKRTA
DGSEFESPKKKRKVSGGSSGGSTLNIEDEYRLHETSKEPDVSLGSTWLSDFPQAWAETGGMGLAVR
QAPLIIPLKATSTPVSIKQYPMSQEARLGIKPHIQRLLDQGILVPCQSPWNTPLLPVKKPGTNDYRPVQ
DLREVNKRVEDIHPTVPNPYNLLSGLPPSHQWYTVLDLKDAFFCLRLHPTSQPLFAFEWRDPEMGIS
GQLTWTRLPQGFKNSPTLFNEALHRDLADFRIQHPDLILLQYVDDLLLAATSELDCQQGTRALLQTL
GNLGYRASAKKAQICQKQVKYLGYLLKEGQRWLTEARKETVMGQPTPKTPRQLREFLGKAGFCRL
FIPGFAEMAAPLYPLTKPGTLFNWGPDQQKAYQEIKQALLTAPALGLPDLTKPFELFVDEKQGYAKG
VLTQKLGPWRRPVAYLSKKLDPVAAGWPPCLRMVAAIAVLTKDAGKLTMGQPLVILAPHAVEALV
KQPPDRWLSNARMTHYQALLLDTDRVQFGPVVALNPATLLPSGGSKRTADGSEFESPKKKRKVGSG
PAAKRVKLD*

Extended data table 6: List of antibodies used in this study.

| antibody | clone | host species | dilution | application |
|---|---|---|---|---|
| FITC anti-CD11b | M1/70 | rat | 1:50 | FACS |
| PE Biotin | Bio3-18E7 | mouse | 1:50 | FACS |
| PE-Vio®615 ACSA-2 | REA969 | rec-human | 1:50 | FACS |
| APC-O4 | REA576 | rec-human | 1:50 | FACS |
| NEUN | ab104224 | mouse | 1:500 | histology |
| GFAP | ab95231 | chicken | 1:1'500 | histology |
| *Sp*Cas9 | D8Y4K | rabbit | 1:1'000 | histology |
| anti-rabbit A488 | JIR-711-545-152 | donkey | 1:1'000 | histology, secondary |
| anti-chicken Cy5 | JIR-703-175-155 | donkey | 1:500 | histology, secondary |

| Extended data table 6 continued. | | | | |
|---|---|---|---|---|
| antibody | clone | host species | dilution | application |
| anti-mouse Cy3 | JIR-715-165-151 | donkey | 1:500 | histology, secondary |
| ADRB1 | ab85037 | rabbit | 1:1'000 | histology, Western blot |
| ACTB | ab8226 | mouse | 1:2'000 | Western blot |

Extended data table 7: Oligos used for deep sequencing.

| oligo name | oligo sequence (5'→ 3') |
|---|---|
| HTS-Adrb1-endogenous_fwd | CTTTCCCTACACGACGCTCTTCCGATCTNNNNNNCCAGCATTGAGACCCTGTGT |
| HTS-Adrb1-endogenous_rev | GGAGTTCAGACGTGTGCTCTTCCGATCTNNNNNNCATGAGGATGGGCAGGAAGG |
| HTS-Adrb1-PiggyBac_rev | GGAGTTCAGACGTGTGCTCTTCCGATCTNNNNNNATAGGGCCCTCTAGACGCTT |
| HTS-Dnmt1-invivo_fwd | CTTTCCCTACACGACGCTCTTCCGATCTNNNNNNGTCTTCCCCCACTCTCTTGC |
| HTS-Dnmt1-invivo_rev | GGAGTTCAGACGTGTGCTCTTCCGATCTNNNNNNCCCCAATATATGCCTCGGC |
| HTS-Adrb1-invivo_fwd | CTTTCCCTACACGACGCTCTTCCGATCTNNNNNNTCGCTACCAGAGTTTGCTGA |
| HTS-Adrb1-invivo_rev | GGAGTTCAGACGTGTGCTCTTCCGATCTNNNNNNAGCACTTGGGGTCGTTGTAG |
| HTS-eIF2B4-PiggyBac_fwd | CTTTCCCTACACGACGCTCTTCCGATCTNNNNNNGTGCTGGAATTCCTAAAAGG |
| HTS-eIF2B4-PiggyBac_rev | GGAGTTCAGACGTGTGCTCTTCCGATCTNNNNNNGTGATCACCAGATCCACGAG |
| HTS-eIF2B5-PiggyBac_fwd | CTTTCCCTACACGACGCTCTTCCGATCTNNNNNNGATCACAGAGTTGGGCAGAC |
| HTS-eIF2B5-PiggyBac_rev | GGAGTTCAGACGTGTGCTCTTCCGATCTNNNNNNCCTTGGGTCTTCTGGAAGTG |
| HTS-Otc-PiggyBac_fwd | CTTTCCCTACACGACGCTCTTCCGATCTNNNNNNGGGAGGACACCCTTCCTTTC |
| HTS-Otc-PiggyBac_rev | GGAGTTCAGACGTGTGCTCTTCCGATCTNNNNNNCAGTCCCTACCTGTGCCCAC |

Extended data table 8: Nucleotide sequences of HTS amplicons.

| amplicon name | amplicon sequence (5'→ 3') |
|---|---|
| *Adrb1* | CCAGCATTGAGACCCTGTGTGTCATCGCCCTGGACCGCTACCTCGCCATCACGTCGCCCTTTCGCTACCAGAGTTTGCTGACGCGCGCGCGAGCGCGGGCCCTCGTGTGCACAGTGTGGGCCATCTCGGCGTTGGTGTCCTTCCTGCCCATCCTCATG |
| *Dnmt1* | GTCTTCCCCCACTCTCTTGCCCTGTGTGGTACATGCTGCTTCCGCTTGCGCCGCCCCCTCCCAATTGGTTTCCGCGCGCGCGAAAAAGCCGGGGTCTCGTTCAGAGCTGTTCTGTCGTCTGCAACCTGCAAGATGCCAGCGCGAACAGCTCCAGCCCGAGTGCCTGCGCTTGCCTCCCCGGCAGGCTCGCTCCCGGACCATGTCCGCAGGCGGTAGGTGCCACGCAGGGTGGGGGTGAGGGGCGGGACCGATGCCGAGGCATATATTGGGGG |
| *EIF2B4* | GTGCTGGAATTCCTAAAAGGATGTTGACACCCACTTGTCATTTCAGATGATCCCGATGATCTGCAGTGTAAGCGGGGAGACCAGGTGGCCCTGGCTAACTGGCAGAGCCACCCGTCCCTCTGGTTGTTGAATCTGGTCTATGACGTGACTCCACCTGAGCTCGTGGATCTGGTGATCAC |
| *EIF2B5* | GATCACAGAGTTGGGCAGACTAACTGTGCCTCTGGTTCTTAATAGGTTAAGAAGGAAGCTAGAAAAAAATGTCTCTGTGATGACAATGGTCTTCAAAGAGTCGTCACCCAGCCACCCTACACACTGCCATGAGGACAACGTGGTGATGGCTGTGGACAGCGCCACCAACAGGGTTCTTCACTTCCAGAAGACCCAAGG |
| *Otc* | GGGAGGACACCCTTCCTTTCTTACCACACAAGACATTCACTTGGGTGTGAATGAAAGTCTCACAGACACCGCTCAGTTTGTAAAACTTTTCTTCCTTCCAAAGTTTATTTCAAACTCTGATGGGTTAGTTTAAAAGAGAAGATGATGCTTCTCCTTAGATAATGGTCTCCCCGGGTGGGCACAGGTAGGGACTG |
